# Viruses in laboratory *Drosophila* and their impact on host gene expression

**DOI:** 10.1101/2023.07.10.548260

**Authors:** Oumie Kuyateh, Darren J Obbard

**Affiliations:** Institute of Ecology and Evolution, Ashworth Laboratories, Charlotte Auerbach Road, Edinburgh EH9 3FL, UK

**Author notes:** Wellcome Trust Genome Campus, Hinxton, Saffron Walden CB10 1SA (Oumie Kuyateh).

**Keywords:** *Drosophila melanogaster*, virus, immunity, transcriptome, Drosophila C virus, Drosophila A virus, nora virus, galbut virus

## Abstract

*Drosophila melanogaster* has one of the best characterized antiviral immune responses among invertebrates. However, relatively few easily-transmitted natural virus isolates are available, and so many *Drosophila* experiments have been performed using artificial infection routes and artificial host-virus combinations. These may not reflect natural infections, especially for subtle phenotypes such as gene expression. Here, to explore the laboratory virus community and to better understand how natural virus infections induce changes in gene expression, we analyse seven publicly available *D. melanogaster* transcriptomic sequencing libraries that were originally sequenced for projects unrelated to virus infection. We find ten known viruses—including five that have not been experimentally isolated—but no previously unknown viruses. Our analysis of host gene expression found numerous genes were differentially expressed in flies that were naturally infected with a virus. For example, flies infected with nora virus showed patterns of gene expression consistent with intestinal vacuolization and host attempted repair via the upd3 JAK/STAT pathway. We also found marked sex-differences in virus-induced differential gene expression. Our results show that natural virus infection in laboratory *Drosophila* does indeed induce detectable changes in gene expression, suggesting that this may form an important background condition for experimental studies in the laboratory.

## Introduction

The antiviral immune response of *Drosophila melanogaster* is among the best characterised of any invertebrate, and antiviral responses in *Drosophila* are mediated by pathways that are conserved across many taxa (Mussabekova et al., 2017). These include RNA interference (RNAi), the immune deficiency pathway (Imd), the Toll-Dorsal pathway (Toll), Janus kinase/signal transducer, activator of transcription pathway (JAK/STAT), autophagy, and the stimulator of interferon genes (STING) signalling cascade (Dostert et al., 2005; Holleufer et al., 2021; Liu et al., 2022; Palmer et al., 2019; Shelly et al., 2009; X.-H. Wang et al., 2006; Zambon et al., 2005).

Virus infection may be associated with dramatic changes in gene expression. First, infections trigger host signalling cascades that eventually alter the expression of host immune effector molecules. For example, wild-type *D. melanogaster* infected with Drosophila C Virus (DCV) display increased expression of genes such *Spaetzle* (*Spz*) and *Drosomycin* (*Drs*) that encode a cytokine and an antimicrobial peptide respectively, and which are important in the Toll pathway (Kemp et al., 2013). There is also increased expression of the immune-associated genes CG12780, vir-1, and *listericin* (CG9080) (Hedges & Johnson, 2008). Second, the expression of non-immune genes changes as a consequence of infection, either through viral manipulation of the host, or through the consequences of disease. For example, female flies infected with Drosophila Kallithea Nudivirus display decreased expression of chorion proteins as they cease laying eggs (Palmer et al., 2018).

Much of what is known about the *Drosophila* expression response to viral infection has come from transcriptome profiling studies (e.g. Cordes et al., 2013; Merkling et al., 2015; Palmer et al., 2018; Zhu et al., 2013). Most such studies rely on systemic viral infections, established through microinjection (or septic puncture) into the body cavity. However, the infection route of pathogens can substantially affect the outcome of infection, and can trigger different antiviral responses (Gupta, Vasanthakrishnan, et al., 2017; Mondotte et al., 2018; Mondotte & Saleh, 2018). This may in part be because injection bypasses immune responses at the natural site of pathogen entry, such as cuticle, trachea, gut, and reproductive organs. For example, the Toll pathway appears necessary for resistance to oral DCV infection via activation of the transcription factor Dorsal, whereas it is not apparently required in resistance to systemic DCV infection (Ferreira et al., 2014). The route of infection is also important in shaping viral tropism, host response, and pathology. For example, DCV infection is almost 100% lethal when injected, but causes only 10-25% mortality when flies are infected orally—even with very high viral titres (Ferreira et al., 2014; Wong et al., 2016). Remarkably, oral DCV infections at the larval stage are reported to protect flies during reinfection by injection as adults (Mondotte et al., 2018). In contrast, it has been reported that flies injected with sublethal dose of DCV are not protected against subsequent DCV injection, indicating the importance of oral infection in priming the immune response (Longdon et al., 2013).

A relative lack of natural virus isolates may also limit studies of host-virus interaction using the *Drosophila* model. Such interactions can be highly host-specific (Tafesh-Edwards & Eleftherianos, 2020), for example it has been suggested that the Virus protein 1 (VP1) of Drosophila immigrans nora virus is unable to suppress the antiviral RNAi pathway in *D. melanogaster*, whereas it can suppress it in *D. immigrans* (van Mierlo et al., 2014). However, many studies in *Drosophila* have used non-native viruses such as Flock House Virus (a beetle virus, e.g. Dostert et al., (2005)), Sindbis virus (a mosquito alphavirus, e.g. Avadhanula et al., (2009)), Invertebrate Iridescent Virus 6 (a moth iridovirus, e.g. Bronkhorst et al., (2012)) and Cricket Paralysis Virus (a moth dicistrovirus, e.g. Nayak et al., (2010)).

Despite the fact that metagenomic sequencing of wild *Drosophila* has revealed over 120 naturally occurring fruit fly viruses, spanning more than 25 families (Medd et al., 2018; Shi et al., 2018; Wallace et al., 2021; Webster et al., 2015, 2016), until recently relatively few natural *Drosophila melanogaster* viruses had isolates available; only the RNA viruses DCV, *D. melanogaster* Sigmavirus (DmelSV), *Drosophila* A virus (DAV), *Drosophila* X virus (DXV), nora virus, and the DNA virus, Drosophila Kallithea nudivirus virus (Habayeb et al., 2006; Jousset et al., 1972; Palmer et al., 2018; Teninges et al., 1979). More recently, galbut virus, La Jolla virus, and Newfield virus have also been isolated (Bruner-Montero et al., 2023; Cross et al., 2020), but these have not yet been widely used in experimental studies.

Some of these viruses are known to occur in laboratory flies and cell culture. The most common viruses reported from laboratory fly stocks include DCV, nora virus, DAV, Newfield virus, DMelSV, and Thika virus (Webster et al., 2015). In addition, Drosophila X virus, Drosophila Totivirus, Newfield virus, American Nodavirus, Bloomfield virus, nora virus, DCV, and DAV have all been found laboratory cell cultures (Echalier, 1997; Webster et al., 2015). The widespread occurrence of such viruses in experimental stocks raises the question of whether changes in gene expression induced by these viruses can impact laboratory experiments. For example, viruses can affect fecundity and shorten development time and lifespan of *Drosophila* (Brosh et al., 2022; Gomariz-Zilber & Thomas-Orillard, 1993; Gupta, Stewart, et al., 2017; Palmer et al., 2018) and can change fruit fly behaviour and mobility (e.g. Brun & Sigot, 1955; Gupta, Stewart, et al., 2017; Palmer et al., 2018), which may negatively impact the interpretability of life-history and developmental studies.

Here we survey seven publicly-available adult transcriptome project datasets from laboratory *D. melanogaster* to quantify the prevalence of viruses in experimental studies, and to assess the impact of viruses on patterns of host gene expression. As these infections were incidental and unintended by the original authors, they reflect natural laboratory infection routes and host virus combinations. The resulting changes in gene expression are suggestive of an reduction in host reproductive investment, and nora virus induced gut pathology and host repair. These results imply that background changes in gene expression due to viral infection may be relevant to laboratory experiments.

## Materials and Methods

### Data sources

We initially selected nine *D. melanogaster* RNAseq ‘projects’ that each comprised at least 130 sequencing libraries, and downloaded them from the European Nucleotide Archive (*European Nucleotide Archive*, 2022, **Table 1**). The selected datasets reflected a range of original experimental purposes. For example, an exploration of natural genetic variation in expression regulation in the Drosophila Genetic Reference Panel (DGRP) (Everett et al., 2020; Lin et al., 2016), the relationship between genotype and circadian gene expression (Litovchenko et al., 2021), the utility of different bioinformatics pipelines (Everett et al., 2020), the impact of lead exposure on expression across development (Qu et al., 2017), and the role of selection in sex-biased gene expression (Ingleby et al., 2016). Two projects lacked any evidence of viral infection (PRJNA527284 and PRJNA52728), and these were excluded. Two projects comprised a mix of developmental stages and/or cell culture (PRJNA305983 and PRJNA75285), and from these we retained only adult data.

**Table 1.**
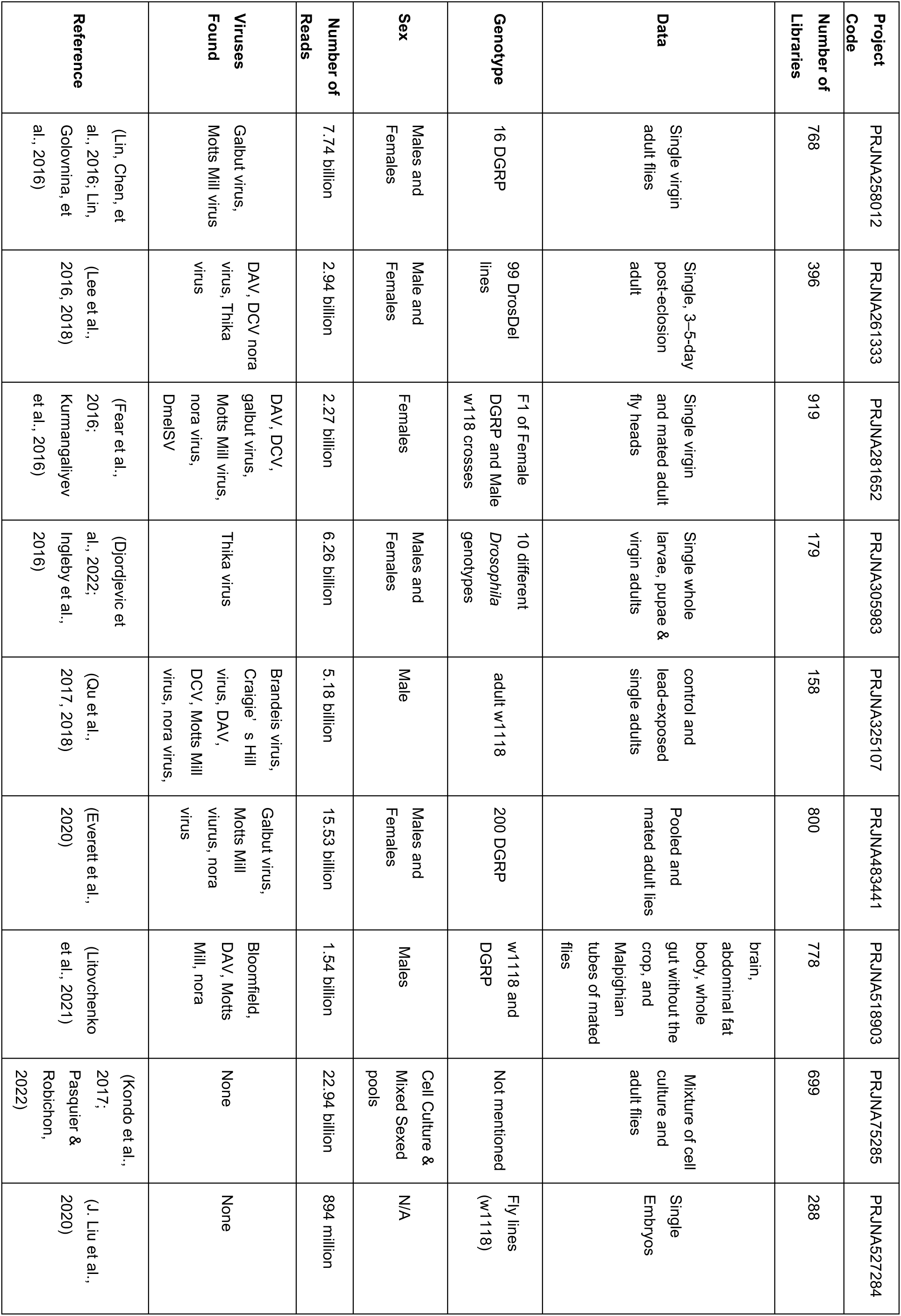
Datasets examined

### Virus detection and quantification of host expression

To identify unknown and potentially novel viral infections, we *de novo* assembled non-fly reads. We first excluded any reads mapping to *D. melanogaster* or to well-studied contaminating cellular organisms such as bacteria, fungi, trypanosomatids, and microsporidia (a ‘hologenome’ as reported in Wallace et al., 2021) using Bowtie 2 (Langmead & Salzberg, 2012). We then assembled the remaining read pairs using Trinity (Grabherr et al., 2011), and retained all scaffolds with a length of at least 500 nt. We grouped the resulting scaffolds into clusters meeting at least a 95% sequence identity threshold using CD-HIT (Fu et al., 2012). Cluster representatives were then used to search against a custom database using Diamond blastx (Buchfink et al., 2014), retaining clusters with a score at most 5% lower than the best alignment score for each query. This custom target database comprised all viral proteins from the Genbank non-redundant (nr) protein database (Clark et al., 2016) and all the prokaryote, protist, fungal, nematode, hymenopteran, and dipteran sequences from NCBI refseq protein database (O’Leary et al., 2016). Contigs from each virus were then assembled together using Geneious Prime. These manually curated viral references were then used as targets for viral quantification by mapping using Bowtie 2 (Langmead & Salzberg, 2012). To mitigate the potential impact of barcode-switching (Griffiths et al., 2018), the a virus was considered to be present in a library if the number of reads from that virus was at least 1% of the number in the library with the highest number of reads from that virus. We additionally applied a minimum threshold of 150 reads, chosen as this threshold reduced the inconsistency between duplicate samples (**Figure S1**). For comparison, we also estimated virus presence/absence in each library by estimating the rate of barcode switching based on sex-specific genes, but this approach gave similar results (**Document S1**).

To quantify fly gene expression, we mapped reads as pairs to the *D. melanogaster* genome (FlyBase release r6.27, FEB2019, ‘FlyBase’, 1996) using the splice-aware mapper, STAR (Dobin et al., 2013) and counted the mapped reads using ‘featureCounts’ (Liao et al., 2014). We verified the reported sex of each fly by counting reads mapping to twelve male-specific and five female-specific genes (FBgn0011669, FBgn0011694, FBgn0046294, FBgn0053340, FBgn0250832, FBgn0259795, FBgn0259971, FBgn0259975, FBgn0262099, FBgn0262623, and FBgn0270925 versus FBgn0000355, FBgn0000356, FBgn0000358, FBgn0000359, FBgn0000360, respectively. All gene counts from featureCounts and metadata including the number of reads that mapped to each virus per dataset are presented in (**Zip File S1**)

### Virus prevalence and diversity

We quantified virus prevalence as the proportion of sequencing libraries within each published project for which the virus read number passed our presence/absence threshold (selected based on the inferred barcode switching rate, above). We then assumed binomial sampling to obtain confidence intervals for the proportion of affected libraries in each project. To explore the relationships between laboratory viruses and previously published virus sequences, we selected the sample within each project that had the highest virus read number for each virus. For galbut virus, which has two clearly distinct haplotypes, we selected the sample with the highest read number for each haplotype. We then performed phylogenetic analyses using the sequences from these samples, and incorporating all the other sequences from each virus available in GenBank (Clark et al., 2016). For analysis of segmented viruses, we selected the segment that had the most examples in genbank. The number of sequences available varied between 3 (Brandeis virus) and 144 (DMelSV), and the aligned sequence length varied between 1.4kb (galbut virus) and 11.3kb (nora virus). DNA sequences were aligned using MAFFT (Katoh, 2002) and we inferred maximum likelihood trees using iqtree version 2 (Minh et al., 2020), using the best supported substitution model according to the Bayesian information criterion. Note that recombination is common in many of these viruses (Webster et al., 2015), such that trees should be interpreted in terms of overall similarity rather than relationship, and that branch lengths will not be proportional to divergence time. All alignments, model details, and trees are presented in (**Zip File S2**)

### Differential host gene expression

We performed a differential gene expression analyses using ‘Dream’ (Hoffman & Roussos, 2021), an R package (R Core Team, 2021) for gene expression analysis in R that permits the use of mixed effect models. Each virus was analysed separately by combining all the project datasets in which it was detected, treating the presence/absence of each virus in each library as a fixed effect in the differential expression model. Host sex was also fitted as a fixed effect, as was its interaction with the presence/absence of the virus. The genotype and dataset project codes were combined and fitted as a random effect to account for differences between genetic background and laboratory environment. If different treatment methods, mating statuses, or tissues were reported, these were also combined and included in the random effect. If both sexes were present, uninfected females were treated as the reference condition for comparisons. For example, if two viruses were present in a study, and both sexes were sampled from multiple genotypes and tissues, our analysis model would be:

𝐸𝑥𝑝𝑟𝑒𝑠𝑠𝑖𝑜𝑛 ∼ 𝑆𝑒𝑥 + 𝑉𝑖𝑟𝑢𝑠 + 𝑆𝑒𝑥: 𝑉𝑖𝑟𝑢𝑠 + (1|𝐷𝑎𝑡𝑎𝑠𝑒𝑡_𝐺𝑒𝑛𝑜𝑡𝑦𝑝𝑒_𝑇𝑖𝑠𝑠𝑢𝑒)

If the virus was present in only one sex in a mixed-sex project dataset, or the infected project datasets had only one sex, then the sex term and its interaction with virus were excluded. If a virus was present in only one dataset then the dataset term was excluded from the model, and if the dataset had only one genotype, then the genotype term was also removed accordingly. A cut-off log-fold change logFC > 0.5 and logFC < -0.5 were used to define increased and decreased expression respectively, and an Benjamini-hochberg adjusted p-value < 0.001 (to account for multiple testing) was used to indicate statistical significance. The genes with significant changes in expression from the differential expression analysis were subsequently analysed for Gene Ontology enrichment using VISEAGO (Brionne et al., 2019), which depends on TOPGO (Alexa & Rahnenfuhrer, 2016).

### Correlation analysis

To compare the effects on the host by viral infection *post hoc*, we calculated a pearson correlation matrix between the estimated changes in gene expression in response to viruses and their interactions with sex using the rcorr function in the Hmisc package (Harrel Jr, 2022) in R (R Core Team, 2021).

## Results

### Ten RNA viruses were detectable in laboratory RNAseq datasets

We examined a total of nine publicly available RNAseq projects, but two of these contained no viral reads and were excluded from further analyses (**Table 1**). One project (PRJNA258012) contained reads from Flock House virus, Newfield virus, Drosophila Totivirus, and Drosophila X Virus (DXV). All of these viruses are common cell culture contaminants (Webster et al., 2015), and three have previously only been reported from cell culture. In addition, read numbers were very highly correlated among these viruses across sequencing libraries (r≥0.96, p-value=2.2e-16), consistent with a common source (**Figure S2**). We therefore chose to exclude these viruses from further analysis, as we believe they are likely to represent RNA contamination or barcode-hopping from unreported cell culture libraries, rather than infections of the sequenced flies. Neither of the remaining two viruses seen in this project have been reported from cell culture. After excluding these putatively contaminating reads, we detected a total of ten viruses with a high level of confidence, i.e. at high copy number in multiple libraries. These included galbut virus (*Partitiviridae*), Motts Mill virus (*Solemoviridae*), DAV (*Permutotetraviridae*), DCV *(Dicistroviridae),* nora virus (unclassified *Picornovirales*), Dmel Sigma virus (Rhabdoviridae), Bloomfield virus (*Reoviridae*), Craigie’s Hill virus (*Nodaviridae*), Thika virus (unclassified Picornavirales), and Brandeis virus (cf. Negevirus). Some samples displayed an extremely high virus titre (**Zip File S1**). Most extreme was library SRR3654766 (heads from lead-treated males), in which ca. 70% of non-ribosomal RNA reads derived from DCV. Also at an occasionally very high level were Nora virus and DAV, which constituted 46% of non-ribosomal reads in SRR8522465 and 25% in SRR8522473, respectively (adult guts from DGRP lines 309 and 359). The other viruses generally had a much lower copy-number, with Motts Mill virus reaching a maximum of 4.8% of non-ribosomal RNA reads (SRR7620105; adult males from DGRP line 93), Brandeis virus 3.8% (SRR3654657; heads from adult males), Thika virus 2.3% (SRR1577470; adult male Df(2L)ED247/+ flies), and Bloomfield virus 2.4% (SRR8522432; adult guts from DGRP line 380).

Of these ten viruses, nora virus had the highest prevalence, detectable in five projects with a prevalence of up to 53% (PRJNA261333; **Figure 1**). In PRJNA518903, 100% of the nora virus infected libraries were gut tissue—consistent with its known gut tropism (Habayeb et al., 2009). DAV had the second highest prevalence, occurring in four projects with a prevalence of up to 17% (PRJNA281652). The rarest viruses were Craigie’s Hill virus, Bloomfield virus, DMelSV and Brandeis virus, each appearing in one project at prevalences as low as 0.3% (Bloomfield virus in PRJNA518903; **Figure 1**). The two viruses that are thought to be exclusively vertically transmitted (DMelSV and galbut virus; (Contamine & Gaumer, 2008; Cross et al., 2020) were only found in DGRP lines, consistent with those lines’ relatively recent wild origin. And, consistent with previous studies of galbut virus (Webster et al., 2015), we detected two distinct strains with pairwise sequence identity of 85% in DGRP projects PRJNA258012 and PRJNA483441. It is noteworthy that no laboratory isolates have been published for Thika virus, Motts Mill virus, Brandeis virus, Craigie’s Hill virus, and Bloomfield virus, potentially allowing us to analyse *Drosophila* expression in response to these pathogens for the first time.

**Figure 1.**
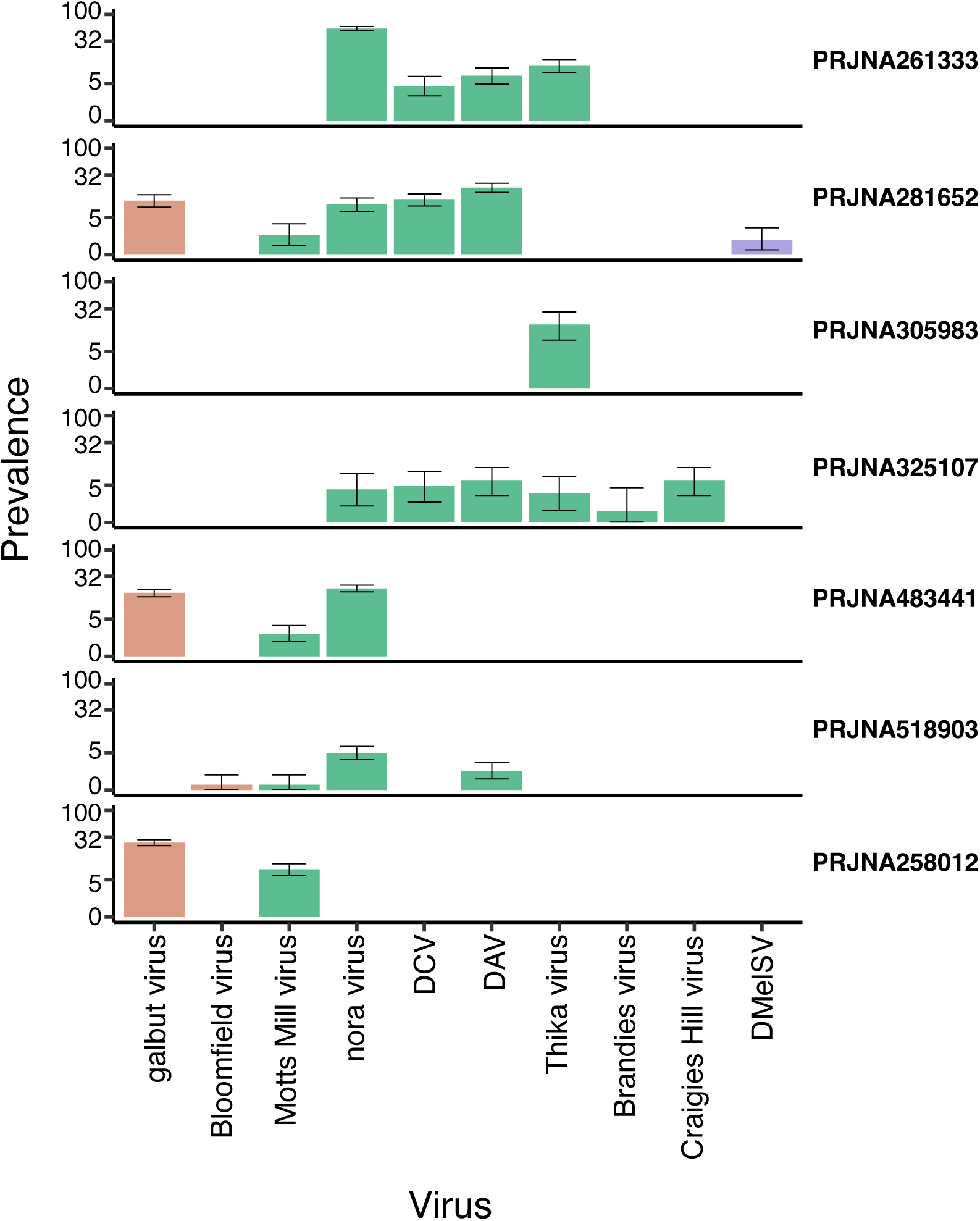
Prevalence of Viruses in the laboratory *D. melanogaster*. The figure shows the prevalence of galbut virus, Motts Mill virus, DAV, DCV, nora virus, DmelSV, Bloomfield virus, Craigie’s Hill virus, Thika virus, and Brandeis virus in the seven projects analysed. The dataset project codes are shown for each, and brown, green, and mauve bar charts represent dsRNA, +ssRNA, and -ssRNA viruses. Prevalence is given as a percentage of libraries in which the virus was detected, plotted on a log scaled axis for clarity, with 95% confidence intervals assuming binomial sampling.

### Laboratory viruses are closely related to each other

Our phylogenetic analyses showed that although laboratory viruses were very rarely identical across projects, viruses from different projects sometimes clustered together with each other and previously published laboratory isolates. This was most notable for DAV, DCV, and Nora virus (A-C; **Figure S3**), and may suggest that there are clades of these viruses circulating in the laboratory environment. The same may be true for Motts Mill virus, as the four sequences formed two near-identical pairs, but previous laboratory isolates have not been reported (**Document S2)**

### Many laboratory virus infections did not induce detectable changes in gene expression

We did not detect any significant changes in gene expression in response to DMelSV (**Spreadsheet S1**; **Figure S4**), and most of the expression changes in response to Bloomfield virus (**Figure S4**), galbut virus, and DCV were similarly not significant (**Figure 2**). *Tret1-2*, which encodes a sugar transporter (Kanamori et al., 2010), was the only gene with a significant change in expression in response to DCV, and *dpr6*, *CG4676*, and *Apl* were the only genes with significant increase in response to galbut virus infection . *Defective proboscis extension response 6* (*dpr6*) is involved in synapse organisation (Carrillo et al., 2015) whilst *CG4676* and *Apl* are involved in protein transport and localisation (Bannan et al., 2008; Boeynaems et al., 2016). In Bloomfield virus infected flies, only three genes, *Ets21C*, *CG16995*, and *CG18649* were significantly upregulated (**Figure S4**). *Ets at 21C* is involved in response to stress such as infection or oncogene activation, CG16995 is involved in sexual reproduction whilst the function of CG18649 is still unknown.

**Figure 2.**
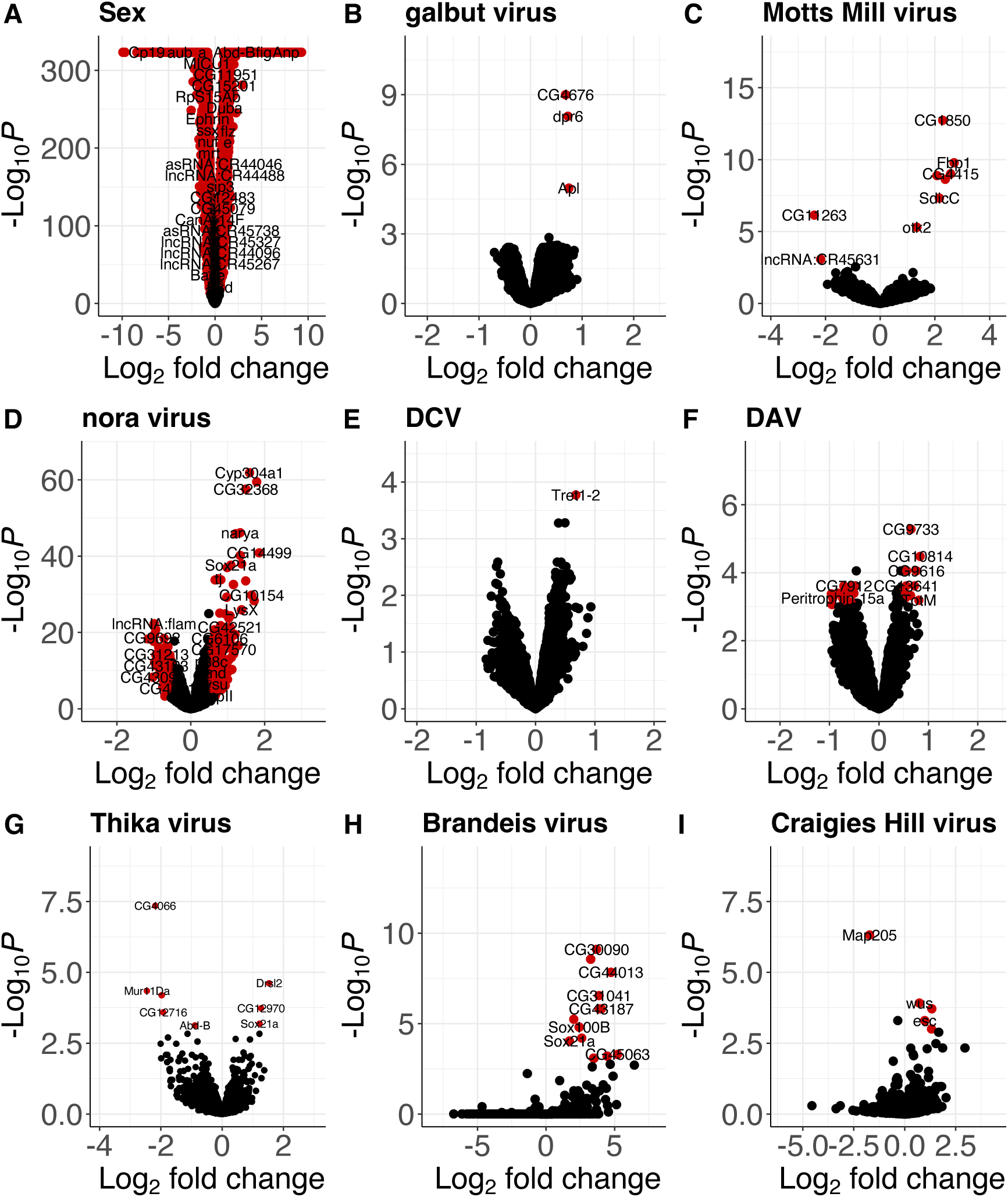
Effects of viral infection on host gene expression. Using uninfected females as baseline, the volcano plots show the main effects on *Drosophila* gene expression of sex (i.e. being male) in panel A, and viral infection in panels B to I. The virus names and total number of variables (genes) used in each expression analysis is shown. A significance threshold of p < 0.001 was used and a logFC cutoff of 0.5. The genes in red are both nominally significant and have a 0.5 < logFC < -0.5.

### Motts Mill virus and Craigies Hill virus affected the expression of genes related to development

Of the 454 genes that displayed a logFC of more than 0.5 in response to Craigies Hill virus infection, only four were nominally significant (adjusted p-value threshold of 0.001; **Figure 2**). The significantly upregulated genes in response to Craigies Hill virus infection include *esc* and *wus* that are involved in development (Lollies, 2012; L. Wang et al., 2006) and *CG31704* that is involved in sexual reproduction (Findlay et al., 2008). *Microtubule-associated protein 205* (*Map-205*) that is involved in mitosis (Dobbelaere et al., 2008), was the only significantly downregulated gene in response to Craigies Hill virus infection.

Of the 1865 genes that showed displayed a logFC of more than 0.5 in response to Motts Mill virus infection, only seven were nominally significant (adjusted p-value threshold of 0.001; **Figure 2**). The upregulated genes in response to Motts Mill virus infection include *Lsp1alpha* and *Fbp1* that encode macromolecular components (Burmester et al., 1999; Wolfe et al., 1977), *SdicC* that is involved in microtubule transport, and *tipE* that enhances para sodium ion channel function and is required during pupal development to rescue adult paralysis (Feng et al., 1995). Of the 3275 genes that showed displayed a logFC of less than 0.5 in response to Motts Mill virus infection, only two were nominally significant (adjusted p-value threshold of 0.001; **Figure 2**). These two downregulated genes were lncRNA:CR45631 and CG11263, which encode a long non-coding RNA and an RNA binding protein.

### Brandeis virus and Thika virus affected the expression of genes related to reproduction and immunity

Out of 1082 genes that showed a logFC of more than 0.5 in response to Brandeis virus, fourteen were nominally significant (adjusted p-value threshold of 0.001; **Figure 2**). These included CG31704 and CG3349 that are involved in male sexual reproduction (Findlay et al., 2008; Sitnik et al., 2016). Other significantly upregulated genes in response to Brandeis virus include SPH93 and *Ir56b* that take part in host antimicrobial defence (Shukla et al., 2019) and response to chemical stimuli respectively (Dweck et al., 2022). There were no significantly downregulated genes in response to Brandeis virus.

Out of 240 genes that showed a logFC of more than 0.5 in response to Thika virus infection, three were nominally significant (p-value threshold of 0.001; **Figure 2**).These were cGAS-like receptor 1 that is involved in STING antiviral response (Holleufer et al., 2021) and *Drsl2* which encodes a peptide with homology to the antifungal peptide encoded by *Drs (Tian et al., 2008)*. Another gene, *Sox21a*, which is involved in stem cell differentiation in the midgut, was also significantly upregulated in response to Thika virus infection (Meng & Biteau, 2015). Out of the 394 genes that displayed a logFC of more than -0.5 (adjusted p-value threshold of 0.001; **Figure 2**) in response to Thika virus infection, five were nominally significant. Most of the significantly downregulated genes in response to Thika virus infection are involved in sexual reproduction. These included chorion proteins such as *CG4066*, *CG12716*, and *Mur11Da* (Tootle et al., 2011), and *Abd-B* that is involved in regulating post-mating responses in females (Gligorov et al., 2013).

### The enteric viruses nora virus and DAV may trigger host innate immune response and gut epithelium repair

Of the 340 genes that showed a logFC of more than 0.5 in response to nora virus infection, 330 were significant (adjusted p-value threshold of 0.001; **Figure 2**). Many of the significantly upregulated genes in response to nora virus infection are involved in midgut stem differentiation. These include *Ser12*, which is predicted encode a protein implicated in wound healing (Ross et al., 2003), and *Ptx1*, *Sox21a*, and *GATAe*, which encode proteins implicated in differentiation in the midgut stem cell (Meng & Biteau, 2015; Okumura et al., 2016; Vorbrüggen et al., 1997). *Unpaired3* (*upd3*), which is involved in gut epithelium repair via the JAK/STAT pathway was also significantly upregulated in response to nora virus infection. Numerous immune response genes were significantly upregulated in response to nora virus infection. These include *Nazo*, which encodes an antiviral effector protein that is expressed downstream of Sting and relish antiviral response (Zhang et al., 2020), *DptA* and *DptB*, which both encode antimicrobial peptides regulated by the ImD pathway (J. H. Lee et al., 2001; Lemaitre et al., 1997), and *Mtk*, which encodes an antifungal peptide regulated by the ImD and Toll pathways (Levashina et al., 1995).

Of the 314 genes that showed a logFC of less than -0.5 in response to nora virus infection, 313 were significant (adjusted p-value threshold of 0.001; **Figure 2**). These significantly downregulated genes include numerous genes that encode constituents of structural constituent of chitin-based larval cuticle such as *Lcp65Ac* and *TwdlR, and TwdlS* (Cornman, 2009; Karouzou et al., 2007; Zuber et al., 2020). There was also a downregulation of genes involved in mitochondrial function such as *Mics1* and *fzo* that encodes proteins that enhance oxidative phosphorylation and enable fusion of the mitochondrion during spermatid differentiation respectively (Hales & Fuller, 1997; H. Meng et al., 2017).

Of the 43 genes that showed a logFC of more than 0.5 in response to DAV, 12 were significant (adjusted p-value threshold of 0.001; **Figure 2**). Antimicrobial genes such as *Srg1*, *CG6429*, and *TotM* were significantly upregulated in response to DAV infection. *Srg1* is involved in STING antiviral signalling whilst *CG6429* and *TotM* are predicted to be involved in immune response (De Gregorio, 2002; Zhong et al., 2013). Of the 397 genes that showed a logFC decrease of more than -0.5 in response to DAV infection, 25 were significant (adjusted p-value threshold of 0.001; **Figure 2**). The majority of the significantly downregulated genes in response to DAV were long non-coding RNA such as *lncRNA:CR45910* and *lncRNA:CR44953* with unknown function. In response to DAV, many genes predicted to be involved in fatty acid-CoA metabolism such as *CG31091* (Subramanian et al., 2013), *CG6300* (Watkins et al., 2007), and *Traf-like* (K.-A. Lee et al., 2018) were significantly downregulated. We also observed a significant downregulation of *Peritrophin-15a* and *E(spl)malpha-BFM*, which are involved in chitin binding and development (Wijffels et al., 2001) and cell fate specification and sensory organ development via Notch signaling (Lu & Li, 2015).

### Sex Virus Interaction

The interaction between virus and sex could only be inferred for viruses that were present in both sexes, namely galbut virus, Motts Mill virus, nora virus, DCV, DAV and Thika virus (**Spreadsheet S2**; **Figure 3**). The effect of sex on the host response to DCV was not significant for any genes in our analysis. However, in the flies infected with DAV, Thika virus, Motts Mill virus, galbut virus, and nora virus, the effect of sex on virus infection induced significant changes in the expression of 1, 4, 83, 95, and 1504 genes respectively (adjusted p-value threshold of 0.001; **Figure 3**). Only *MFS9*, which is predicted to be involved in transmembrane anion transport (Voßfeldt et al., 2012), was significantly downregulated in males infected with DAV. Four genes, *Mur11Da*, *CG4066*, *CG31661*, and *CG6508,* were significantly upregulated in Thika virus infected males. *Mucin related 11Da, CG4066,* and *CG31661* are predicted to be involved in chorion eggshell assembly (Tootle et al., 2011) and *CG6508* is predicted to be involved in cell death and proteolysis (‘FlyBase’, 1996).

**Figure 3.**
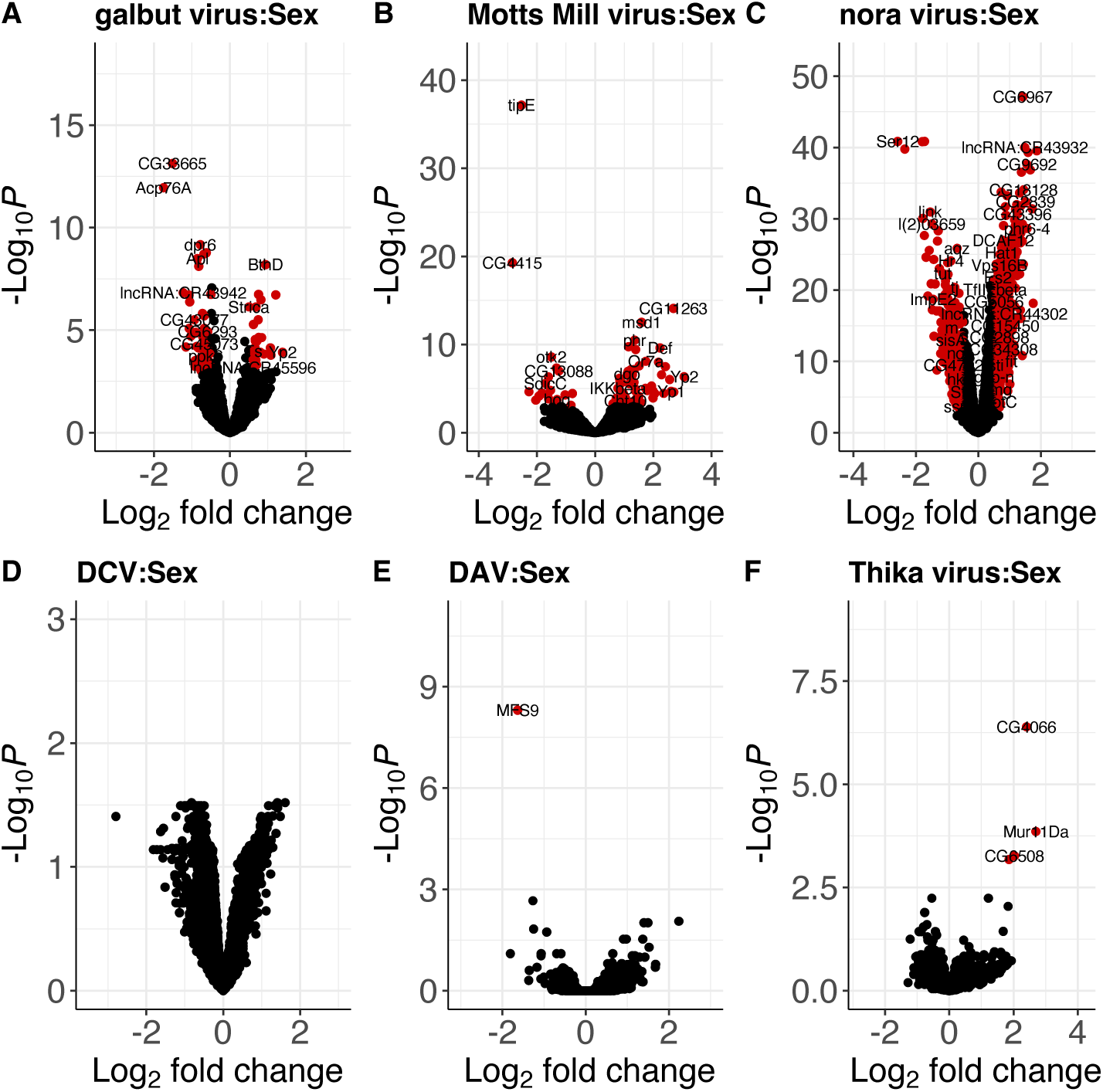
Sex virus Interaction effects on host gene expression. Using uninfected females as baseline, the volcano plots show the sex interaction with viral infection (**A** to **E**), that is the difference in expression between virus-infected males and the expression that would be predicted from the combined effects of sex and infection alone. The virus names and total number of variables (genes) used in each expression analysis is shown. A significance threshold of p < 0.001 was used and a logFC cutoff of 0.5. The genes in red are both significant and have a 0.5 < logFC < - 0.5.

In galbut virus infected males, there was a significant upregulation of numerous genes involved in cuticle development such as *Lcp4* and *Cpr65Ec (Karouzou et al., 2007)* and immune response such as *AttC,* which shows homology to several antimicrobial peptides (Hedengren et al., 2000). In galbut virus infected males, there was also a significant downregulation of male reproductive genes such as *Sfp24Ba* and *Acp76A (Findlay et al., 2008)*,metal ion transport genes such as *dpr3 (Vig et al., 2006)*, and several long non-coding RNA genes (**Figure 3**). In Motts Mill virus infected males, there was a significant upregulation of genes involved in the JAK/STAT pathway such as *GILT2* and *GILT3* (Kongton et al., 2014) and Imd pathway such as *Def and IKKbeta* ((Ertürk-Hasdemir et al., 2009; Verleyen et al., 2006); **Figure 3**). There was also a significant upregulation in the expression of *tipE*, which encodes a protein that enhances sodium ion channel function (Feng et al., 1995) and a significant downregulation of genes implicated in development such as *otk2* and *lov* in Motts Mill virus infected males (Linnemannstöns et al., 2014; Zhou et al., 2016).

In nora virus infected males, there was a significant upregulation of genes such as *Mur82C* and *Cpr56F* that are predicted to encode structural components of extracellular matrix ((Lerch et al., 2020; Syed et al., 2008); **Figure 3**). There was also a significant upregulation of genes involved in systemic immune response such as such as *CecA1* and *Def* that encodes an antimicrobial peptide regulated by the *ImD* and Toll pathways (Verleyen et al., 2006) and male reproductive genes such as *Sfp84E* and *Sfp33A2* (Findlay et al., 2008; Lung & Wolfner, 2001). Nora virus infected males displayed a significant downregulation of immune genes such as and *Ser12* that is involved in wound healing (Campos et al., 2010) and *Send2* that encodes a protein stored in the seminal receptacle and is predicted to be involved in proteolysis (Schnakenberg et al., 2011). The genes *narya* and *ImpE2* that are involved in DNA repair and embryogenesis respectively were also significantly downregulated in males infected with Motts Mill (Lake et al., 2019; Paine-Saunders et al., 1990).

### Changes in gene expression were positively correlated between flies infected with galbut virus, nora virus, and Motts Mill virus

To find if changes in gene expression in response to virus challenge were consistent across viruses, we compared the inferred changes between each of the viruses (**Figure 4**). The magnitude of such correlations was generally small, with the exception of the expression changes induced by galbut virus, DAV, and Motts Mill virus. Changes in gene expression were significantly positively correlated (r = 0.53, p-value < 0.01) between the DAV, galbut virus, and Motts Mill infection, although none were significant in all three (**Figure 4**). Changes in gene expression were significantly positively correlated (r = 0.53, p-value < 0.01) between the galbut:sex and Motts Mill virus:sex. That is, the effect of sex on virus infection induced similar changes in the expression of 209 genes in galbut virus and Motts Mill virus infected flies, with four genes being nominally significant in both infections. These genes include three long non-coding RNA, lncRNA:CR40465, lncRNA:CR44631, lncRNA:CR44289, and *Yp2*, which encodes the major yolk protein of eggs.

**Figure 4.**
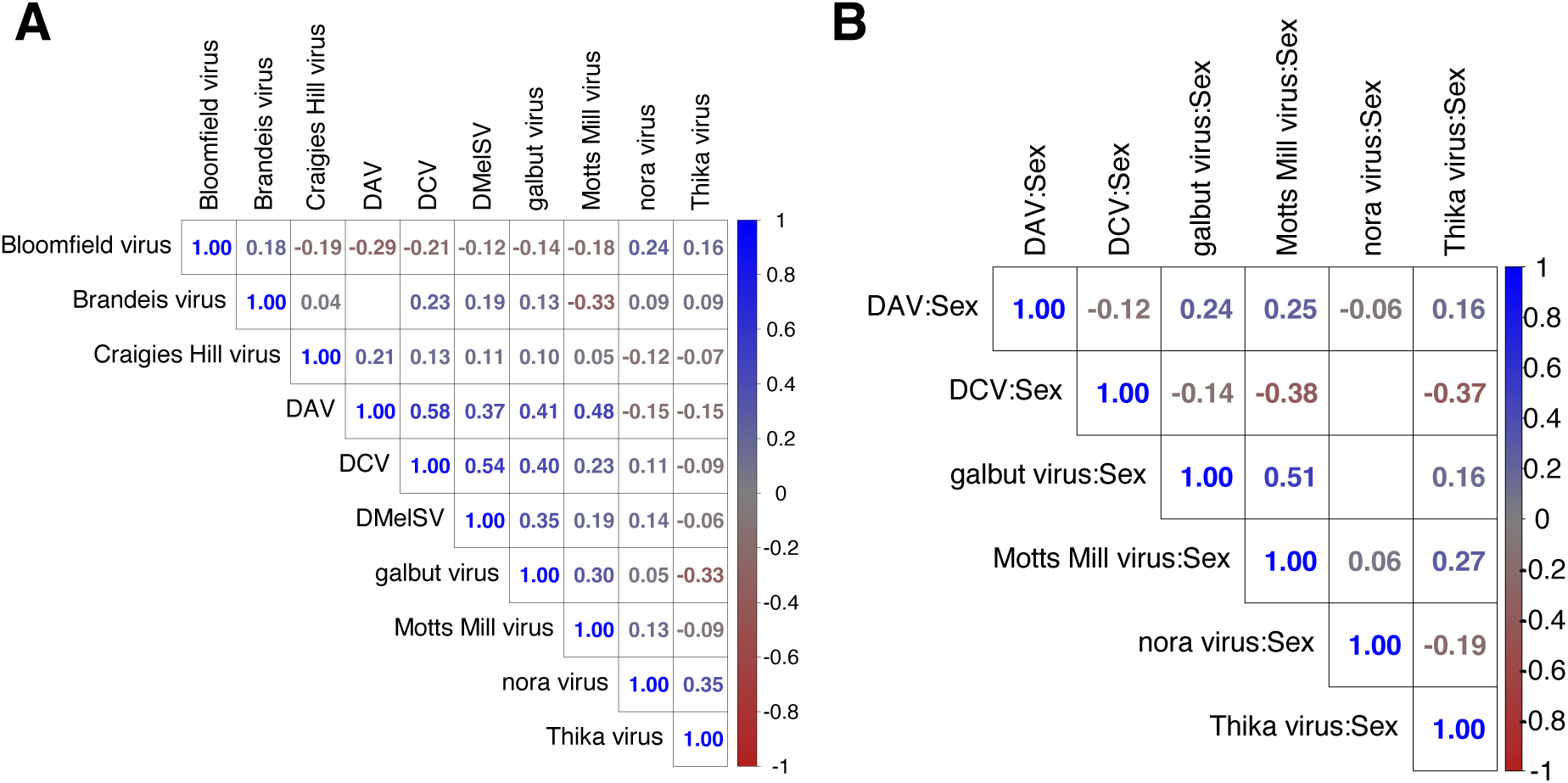
Correlation. The triangular matrix shows significant (p < 0.001) spearman rank correlation coefficients between the estimated changes in gene expression induced by ten *Drosophila* viruses. The virus names are shown above and to the side. Panel **A** shows the correlation between changes in expression in response to viruses. Panel **B** shows the correlations between the changes in expression induced by the effects of sex on viral infection. Blank squares have non-significant correlations.

### Significantly enriched GO Terms

There were no significantly enriched GO terms among the genes that changed expression in response to DmelSV, Bloomfield virus and DCV. Six GO terms related to cell proliferation were significantly enriched in Craigies Hill virus infected flies (**Spreadsheet S3**; **Figure 5**). In general, the enriched GO terms were consistent with the genes that were differentially expressed in our analysis. For example, the significantly enriched terms in response to Thika virus and DAV infection include development and fatty acid metabolism respectively (**Figure 5**). Similar consistency between the significantly enriched GO terms and expression analysis were observed on the effect of sex on virus infection (**Spreadsheet S4**; **Figure 6**). For example, the effect of sex on Motts Mill virus and galbut virus infection significantly enriched GO terms such as ion transport, developmental process, and cuticle development (**Figure 6**). Similarly, the effect of sex on nora virus infection enriched GOP terms related to meiotic cell cycle and response to stimulus (**Figure 6**).

**Figure 5.**
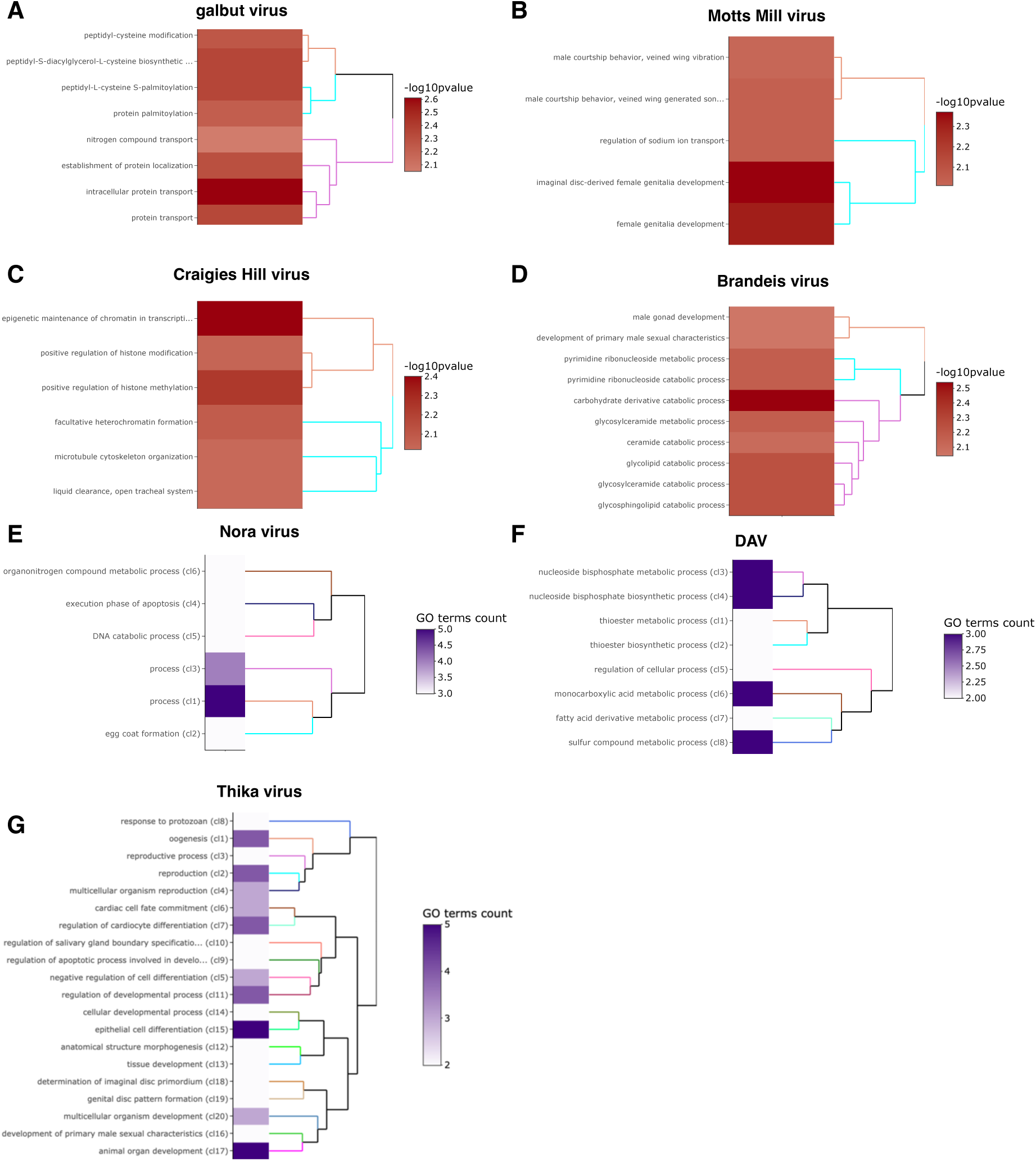
Main Effects significantly enriched GO terms. GO enrichment on the effect of virus infection on *Drosophila* gene expression. The virus names are shown. A, B, C, and D show a -log10(p-value) of functional enrichment analysis and a dendrogram based on hierarchical clustering of the enriched GO terms. The heatmaps of functional GO terms show a clustering combining a description of the first common GO ancestor of each set of GO terms. The heatmap shows the number of GO terms in each set and the dendrogram is based on BMA semantic similarity distance and ward.D2 aggregation criterion (E, F. and G).

**Figure 6.**
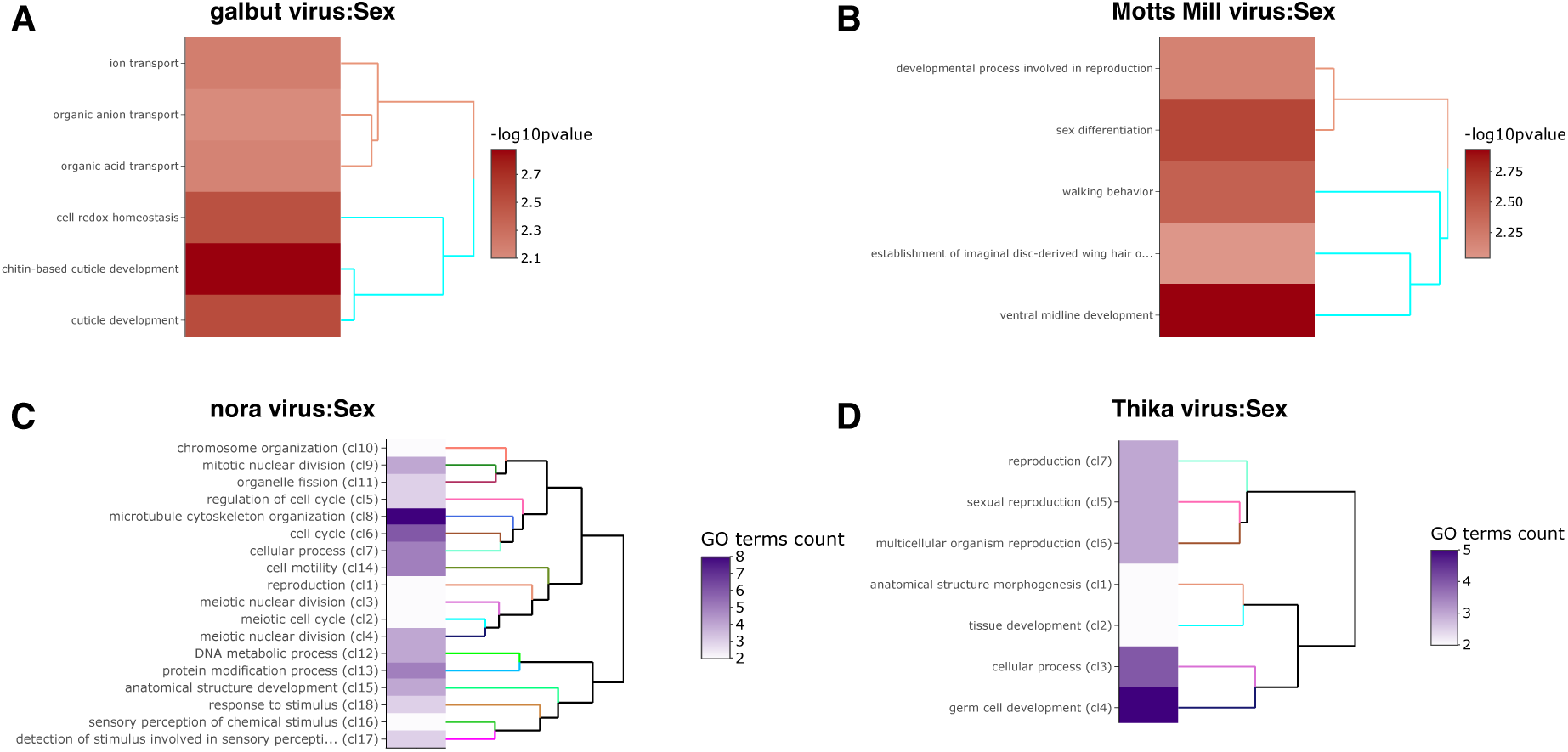
Sex virus interaction significantly enriched GO terms. GO enrichment on the effect of sex on virus infection. The virus names are shown. **A** to **B** show a -log10(p-value) of functional enrichment analysis and a dendrogram based on hierarchical clustering of the enriched GO terms. **C** and **D** are heatmaps of functional GO terms clusters combining a description of the first common GO ancestor of each set of GO terms. The heatmap shows the number of GO terms in each set and the dendrogram is based on BMA semantic similarity distance and ward.D2 aggregation criterion.

## Discussion

Most studies of how *Drosophila* gene expression changes in response to virus infection either use non-native viruses or inject natural viruses into the host—which may not be a good reflection of what happens in nature. To understand how *Drosophila* gene expression changes in response to native virus infection, we analysed seven publicly available RNA-seq datasets that were serendipitously infected with ten different viruses.

### Prevalence

The viruses found in this study were galbut virus, Motts Mill virus, DAV, DCV, nora virus, DmelSV, Bloomfield virus, Craigie’s Hill virus, Thika virus, and Brandeis virus. The most prevalent viruses in the datasets analysed were nora virus and DAV with prevalence of up to 52% and 18% respectively. This is higher than has been observed in wild flies; a study of wild *D. melanogaster* suggested that the average global prevalence of nora virus and DAV is about 7% each (Webster et al., 2015), although the authors also reported that DAV had a prevalence of about 50% in wild *D. melanogaster* collected from Athens (Georgia, USA). The prevalence of galbut virus in this study was 14%, which is much lower than has been reported in wild flies— where the prevalence of galbut is often above 50% (Cross et al., 2020; Webster et al., 2015). The difference in virus prevalence between laboratory and wild flies indicates that some viruses may be better suited than others to the ecology of the lab, suggesting the need to isolate more natural viruses from wild *Drosophila* to better understand host-virus interactions.

### Induction of immune genes in response to viruses

In response to DAV and nora virus infection, we found evidence of increased expression of immune genes involved in antiviral STING pathway, and changes in the expression of genes involved in lipid metabolism. Recent work has shown that flies mutant in STING display reduced lipid storage and downregulated expression of lipid metabolism genes (Akhmetova et al., 2021). The authors also reported that *Drosophila* STING interacts with lipid synthesizing enzymes such as acetyl-CoA carboxylase (Akhmetova et al., 2021).

### Potential virus-induced pathologies

The changes in expression we observe may be indicative of virus induced pathologies on the host. For example, GO analysis showed that changes in the expression of metal ion transport was significantly enriched in the flies infected with Motts Mill virus. Perhaps, virus infection may have induced the deregulation of metal ion transport in flies infected with Motts Mill leading to their change in expression. Conversely, there was an increase in the expression of tipE in response to Motts Mill infection which may have induced changes in the function of the metal ion channels. Temperature-induced paralytic E (tipE) encodes a protein with a cysteine-stabilized αβ scaffold (CSαβ). *Drosophila* peptides with CSαβ scaffold are implicated in the regulation of voltage-gated sodium ion channels (Cohen et al., 2009).

We observed an increase in the expression of genes involved in chitin biogenesis and binding, wound healing, and differentiation in the midgut stem cell respectively. Perhaps, in response to nora virus induced damage of the gut epithelium (Habayeb et al., 2009), the host may activate the JAK/STAT pathway via upd3 in the gut to regulate the repair response. Published work has shown that upd genes are expressed by haemocytes upon gut septic injury, to remotely stimulate stem cell proliferation and the expression of *Drosomycin*-like genes in the intestine (Chakrabarti et al., 2016).

## Funding

Oumie Kuyateh was funded a University of Edinburgh scholarship and the Wellcome Trust PhD Programme in Hosts, Pathogens and Global Health awarded to Professor Keith Matthews (grant 218492/Z/19/Z). For the purpose of open access, the authors have applied a Creative Commons Attribution (CC BY) licence to any Author Accepted Manuscript version arising from this submission.

## Supporting information

Zip File S1

Zip File S2

## Acknowledgments

We thank the original authors of the datasets we analyse for making their data publicly available.

## Author Contributions

Conceptualization, OK and DJO; Methodology, OK.; Writing – Original Draft Preparation, OK.; Writing – Review & Editing, OK and DJO.

## Data Availability Statement

No new data were generated during this study. Analyses and scripts are available from https://github.com/okuyateh/Changes-in-gene-expression-due-to-natural-viral-infection-in-laboratory-Drosophila

## Supplementary Material

**Figure S1:** Best threshold to estimate viral presence.

**Figure S2:** Correlations between normalised reads in PRJNA258012.

**Figure S3:** Phylogenies of nora virus, Thika virus, DAV, and DCV

**Figure S4:** Effects of viral infection on host gene expression.

**Document S1:** Estimating barcode switching rates.

**Document S2:** Phylogenies of virus sequences

**Spreadsheet S1:** Main effects expression data.

**Spreadsheet S2:** Virus sex interaction expression data.

**Spreadsheet S3:** Main effects enriched GO terms.

**Spreadsheet S4:** Virus sex interaction enriched GO terms.

**Zip File S1:** Metadata of datasets and count matrices from featureCounts.

**Zip File S2:** Virus sequence alignments, model details, and trees.

## Notes

https://github.com/okuyateh/Changes-in-gene-expression-due-to-natural-viral-infection-in-laboratory-Drosophila

### Competing Interest Statement

The authors have declared no competing interest.

## Bibliography

1. Akhmetova, K., Balasov, M., & Chesnokov, I. (2021). Drosophila STING protein has a role in lipid metabolism. ELife, 10, e67358. https://doi.org/10.7554/eLife.67358

2. Alexa, A., & Rahnenfuhrer, J. (2016). TopGO: enrichment analysis for Gene Ontology. R Packag. Version 2.26.0. R Package Version 2.26.0.

3. Avadhanula, V., Weasner, B. P., Hardy, G. G., Kumar, J. P., & Hardy, R. W. (2009). A Novel System for the Launch of Alphavirus RNA Synthesis Reveals a Role for the Imd Pathway in Arthropod Antiviral Response. PLoS Pathogens, 5(9), e1000582. https://doi.org/10.1371/journal.ppat.1000582

4. Bannan, B. A., Van Etten, J., Kohler, J. A., Tsoi, Y., Hansen, N. M., Sigmon, S., Fowler, E., Buff, H., Williams, T. S., Ault, J. G., Glaser, R. L., & Korey, C. A. (2008). The Drosophila protein palmitoylome: Characterizing palmitoyl-thioesterases and DHHC palmitoyl-transferases. Fly, 2(4), 198–214. https://doi.org/10.4161/fly.6621

5. Boeynaems, S., Bogaert, E., Michiels, E., Gijselinck, I., Sieben, A., Jovičić, A., De Baets, G., Scheveneels, W., Steyaert, J., Cuijt, I., Verstrepen, K. J., Callaerts, P., Rousseau, F., Schymkowitz, J., Cruts, M., Van Broeckhoven, C., Van Damme, P., Gitler, A. D., Robberecht, W., & Van Den Bosch, L. (2016). Drosophila screen connects nuclear transport genes to DPR pathology in c9ALS/FTD. Scientific Reports, 6(1), 20877. https://doi.org/10.1038/srep20877

6. Brionne, A., Juanchich, A., & Hennequet-Antier, C. (2019). ViSEAGO: A Bioconductor package for clustering biological functions using Gene Ontology and semantic similarity. BioData Mining. https://doi.org/10.1186/s13040-019-0204-1

7. Bronkhorst, A. W., Van Cleef, K. W. R., Vodovar, N., İnce, İ. A., Blanc, H., Vlak, J. M., Saleh, M.-C., & Van Rij, R. P. (2012). The DNA virus Invertebrate iridescent virus 6 is a target of the *Drosophila* RNAi machinery. Proceedings of the National Academy of Sciences, 109(51). https://doi.org/10.1073/pnas.1207213109

8. Brosh, O., Fabian, D. K., Cogni, R., Tolosana, I., Day, J. P., Olivieri, F., Merckx, M., Akilli, N., Szkuta, P., & Jiggins, F. M. (2022). A novel transposable element-mediated mechanism causes antiviral resistance in *Drosophila* through truncating the Veneno protein. Proceedings of the National Academy of Sciences, 119(29), e2122026119. https://doi.org/10.1073/pnas.2122026119

9. Brun, G., & Sigot, A. (1955). Etude de la sensibilité héréditaire au gaz carbonique chez la Drosophile. II. —Installation du virus σ dans la lignée germinale à la suite d’une inoculation. Ann. Inst. Pasteur, 88, 488–513.

10. Bruner-Montero, G., Luque, C. M., Cesar, C. S., Ding, S. D., Day, J. P., & Jiggins, F. M. (2023). Hunting Drosophila viruses from wild populations: A novel isolation approach and characterisation of viruses. PLOS Pathogens, 19(3), e1010883. https://doi.org/10.1371/journal.ppat.1010883

11. Buchfink, B., Xie, C., & Huson, D. H. (2014). Fast and sensitive protein alignment using DIAMOND. In Nature Methods. https://doi.org/10.1038/nmeth.3176

12. Burmester, T., Antoniewski, C., & Lepesant, J.-A. (1999). Ecdysone-regulation of synthesis and processing of Fat Body Protein 1, the larval serum protein receptor of Drosophila melanogaster. European Journal of Biochemistry, 262(1), 49–55. https://doi.org/10.1046/j.1432-1327.1999.00315.x

13. Campos, I., Geiger, J. A., Santos, A. C., Carlos, V., & Jacinto, A. (2010). Genetic Screen in *Drosophila melanogaster* Uncovers a Novel Set of Genes Required for Embryonic Epithelial Repair. Genetics, 184(1), 129–140. https://doi.org/10.1534/genetics.109.110288

14. Carrillo, R. A., Özkan, E., Menon, K. P., Nagarkar-Jaiswal, S., Lee, P.-T., Jeon, M., Birnbaum, M. E., Bellen, H. J., Garcia, K. C., & Zinn, K. (2015). Control of Synaptic Connectivity by a Network of Drosophila IgSF Cell Surface Proteins. Cell, 163(7), 1770–1782. https://doi.org/10.1016/j.cell.2015.11.022

15. Chakrabarti, S., Dudzic, J. P., Li, X., Collas, E. J., Boquete, J.-P., & Lemaitre, B. (2016). Remote Control of Intestinal Stem Cell Activity by Haemocytes in Drosophila. PLOS Genetics, 12(5), e1006089. https://doi.org/10.1371/journal.pgen.1006089

16. Clark, K., Karsch-Mizrachi, I., Lipman, D. J., Ostell, J., & Sayers, E. W. (2016). GenBank. Nucleic Acids Research. https://doi.org/10.1093/nar/gkv1276

17. Cohen, L., Moran, Y., Sharon, A., Segal, D., Gordon, D., & Gurevitz, M. (2009). Drosomycin, an Innate Immunity Peptide of Drosophila melanogaster, Interacts with the Fly Voltage-gated Sodium Channel. Journal of Biological Chemistry, 284(35), 23558–23563. https://doi.org/10.1074/jbc.M109.023358

18. Contamine, D., & Gaumer, S. (2008). Sigma Rhabdoviruses. In Encyclopedia of Virology (pp. 576–581). Elsevier. https://doi.org/10.1016/B978-012374410-4.00503-3

19. Cordes, E. J., Licking-Murray, K. D., & Carlson, K. A. (2013). Differential gene expression related to Nora virus infection of Drosophila melanogaster. Virus Research, 175(2), 95–100. https://doi.org/10.1016/j.virusres.2013.03.021

20. Cornman, R. S. (2009). Molecular Evolution of Drosophila Cuticular Protein Genes. PLoS ONE, 4(12), e8345. https://doi.org/10.1371/journal.pone.0008345

21. Cross, S. T., Maertens, B. L., Dunham, T. J., Rodgers, C. P., Brehm, A. L., Miller, M. R., Williams, A. M., Foy, B. D., & Stenglein, M. D. (2020). Partitiviruses Infecting Drosophila melanogaster and Aedes aegypti Exhibit Efficient Biparental Vertical Transmission. Journal of Virology, 94(20), e01070–20. https://doi.org/10.1128/JVI.01070-20

22. De Gregorio, E. (2002). The Toll and Imd pathways are the major regulators of the immune response in Drosophila. The EMBO Journal, 21(11), 2568–2579. https://doi.org/10.1093/emboj/21.11.2568

23. Dobbelaere, J., Josué, F., Suijkerbuijk, S., Baum, B., Tapon, N., & Raff, J. (2008). A Genome-Wide RNAi Screen to Dissect Centriole Duplication and Centrosome Maturation in Drosophila. PLoS Biology, 6(9), e224. https://doi.org/10.1371/journal.pbio.0060224

24. Dobin, A., Davis, C. A., Schlesinger, F., Drenkow, J., Zaleski, C., Jha, S., Batut, P., Chaisson, M., & Gingeras, T. R. (2013). STAR: Ultrafast universal RNA-seq aligner. Bioinformatics. https://doi.org/10.1093/bioinformatics/bts635

25. Dostert, C., Jouanguy, E., Irving, P., Troxler, L., Galiana-Arnoux, D., Hetru, C., Hoffmann, J. A., & Imler, J.-L. (2005). The Jak-STAT signaling pathway is required but not sufficient for the antiviral response of drosophila. Nature Immunology, 6(9), 946–953. https://doi.org/10.1038/ni1237

26. Dweck, H. K. M., Talross, G. J. S., Luo, Y., Ebrahim, S. A. M., & Carlson, J. R. (2022). Ir56b is an atypical ionotropic receptor that underlies appetitive salt response in Drosophila. Current Biology, 32(8), 1776–1787.e4. https://doi.org/10.1016/j.cub.2022.02.063

27. Echalier, G. (1997). Drosophila viruses and other Infections of Cultured Cells. Drosophila Cells in Culture: Academic Press, 555–595.

28. Ertürk-Hasdemir, D., Broemer, M., Leulier, F., Lane, W. S., Paquette, N., Hwang, D., Kim, C.-H., Stöven, S., Meier, P., & Silverman, N. (2009). Two roles for the *Drosophila* IKK complex in the activation of Relish and the induction of antimicrobial peptide genes. Proceedings of the National Academy of Sciences, 106(24), 9779–9784. https://doi.org/10.1073/pnas.0812022106

29. European Nucleotide Archive. (2022). https://www.ebi.ac.uk/ena/browser/home

30. Everett, L. J., Huang, W., Zhou, S., Carbone, M. A., Lyman, R. F., Arya, G. H., Geisz, M. S., Ma, J., Morgante, F., St. Armour, G., Turlapati, L., Anholt, R. R. H., & Mackay, T. F. C. (2020). Gene expression networks in the *Drosophila* Genetic Reference Panel. Genome Research, 30(3), 485–496. https://doi.org/10.1101/gr.257592.119

31. Feng, G., Deak, P., Chopra, M., & Hall, L. M. (1995). Cloning and functional analysis of tipE, a novel membrane protein that enhances drosophila para sodium channel function. Cell, 82(6), 1001–1011. https://doi.org/10.1016/0092-8674(95)90279-1

32. Ferreira, Á. G., Naylor, H., Esteves, S. S., Pais, I. S., Martins, N. E., & Teixeira, L. (2014). The Toll-Dorsal Pathway Is Required for Resistance to Viral Oral Infection in Drosophila. PLoS Pathogens, 10(12), e1004507. https://doi.org/10.1371/journal.ppat.1004507

33. Findlay, G. D., Yi, X., MacCoss, M. J., & Swanson, W. J. (2008). Proteomics Reveals Novel Drosophila Seminal Fluid Proteins Transferred at Mating. PLoS Biology, 6(7), e178. https://doi.org/10.1371/journal.pbio.0060178

34. FlyBase: The Drosophila database. The Flybase Consortium. (1996). Nucleic Acids Research, 24(1), 53–56. https://doi.org/10.1093/nar/24.1.53

35. Gligorov, D., Sitnik, J. L., Maeda, R. K., Wolfner, M. F., & Karch, F. (2013). A Novel Function for the Hox Gene Abd-B in the Male Accessory Gland Regulates the Long-Term Female Post-Mating Response in Drosophila. PLoS Genetics, 9(3), e1003395. https://doi.org/10.1371/journal.pgen.1003395

36. Gomariz-Zilber, E., & Thomas-Orillard, M. (1993). Drosophila C virus and Drosophila hosts: A good association in various environments. Journal of Evolutionary Biology, 6(5), 677–689. https://doi.org/10.1046/j.1420-9101.1993.6050677.x

37. Grabherr, M. G., Haas, B. J., Yassour, M., Levin, J. Z., Thompson, D. A., Amit, I., Adiconis, X., Fan, L., Raychowdhury, R., Zeng, Q., Chen, Z., Mauceli, E., Hacohen, N., Gnirke, A., Rhind, N., di Palma, F., Birren, B. W., Nusbaum, C., Lindblad-Toh, K., … Regev, A. (2011). Full-length transcriptome assembly from RNA-Seq data without a reference genome. Nature Biotechnology, 29(7), 644–652. https://doi.org/10.1038/nbt.1883

38. Griffiths, J. A., Richard, A. C., Bach, K., Lun, A. T. L., & Marioni, J. C. (2018). Detection and removal of barcode swapping in single-cell RNA-seq data. Nature Communications, 9(1), 2667. https://doi.org/10.1038/s41467-018-05083-x

39. Gupta, V., Stewart, C. O., Rund, S. S. C., Monteith, K., & Vale, P. F. (2017). Costs and benefits of sublethal Drosophila C virus infection. Journal of Evolutionary Biology, 30(7), 1325–1335. https://doi.org/10.1111/jeb.13096

40. Gupta, V., Vasanthakrishnan, R. B., Siva-Jothy, J., Monteith, K. M., Brown, S. P., & Vale, P. F. (2017). The route of infection determines *Wolbachia* antibacterial protection in *Drosophila*. Proceedings of the Royal Society B: Biological Sciences, 284(1856), 20170809. https://doi.org/10.1098/rspb.2017.0809

41. Habayeb, M. S., Cantera, R., Casanova, G., Ekström, J.-O., Albright, S., & Hultmark, D. (2009). The Drosophila Nora virus is an enteric virus, transmitted via feces. Journal of Invertebrate Pathology, 101(1), 29–33. https://doi.org/10.1016/j.jip.2009.02.003

42. Habayeb, M. S., Ekengren, S. K., & Hultmark, D. (2006). Nora virus, a persistent virus in Drosophila, defines a new picorna-like virus family. Journal of General Virology, 87(10), 3045–3051. https://doi.org/10.1099/vir.0.81997-0

43. Hales, K. G., & Fuller, M. T. (1997). Developmentally Regulated Mitochondrial Fusion Mediated by a Conserved, Novel, Predicted GTPase. Cell, 90(1), 121–129. https://doi.org/10.1016/S0092-8674(00)80319-0

44. Hedengren, M., Borge, K., & Hultmark, D. (2000). Expression and Evolution of the Drosophila Attacin/Diptericin Gene Family. Biochemical and Biophysical Research Communications, 279(2), 574–581. https://doi.org/10.1006/bbrc.2000.3988

45. Hedges, L. M., & Johnson, K. N. (2008). Induction of host defence responses by Drosophila C virus. Journal of General Virology, 89(6), 1497–1501. https://doi.org/10.1099/vir.0.83684-0

46. Hoffman, G. E., & Roussos, P. (2021). Dream: Powerful differential expression analysis for repeated measures designs. Bioinformatics, 37(2), 192–201. https://doi.org/10.1093/bioinformatics/btaa687

47. Holleufer, A., Winther, K. G., Gad, H. H., Ai, X., Chen, Y., Li, L., Wei, Z., Deng, H., Liu, J., Frederiksen, N. A., Simonsen, B., Andersen, L. L., Kleigrewe, K., Dalskov, L., Pichlmair, A., Cai, H., Imler, J.-L., & Hartmann, R. (2021). Two cGAS-like receptors induce antiviral immunity in Drosophila. Nature, 597(7874), 114–118. https://doi.org/10.1038/s41586-021-03800-z

48. Ingleby, F. C., Webster, C. L., Pennell, T. M., Flis, I., & Morrow, E. H. (2016). *Sex-biased gene expression in Drosophila melanogaster is constrained by ontogeny and genetic architecture* [Preprint]. Genetics. https://doi.org/10.1101/034728

49. Jousset, F. X., Plus, N., Croizier, G., & Thomas, M. (1972). [Existence in Drosophila of 2 groups of picornavirus with different biological and serological properties]. Comptes Rendus Hebdomadaires Des Seances De l’Academie Des Sciences. Serie D: Sciences Naturelles, 275(25), 3043–3046.

50. Kanamori, Y., Saito, A., Hagiwara-Komoda, Y., Tanaka, D., Mitsumasu, K., Kikuta, S., Watanabe, M., Cornette, R., Kikawada, T., & Okuda, T. (2010). The trehalose transporter 1 gene sequence is conserved in insects and encodes proteins with different kinetic properties involved in trehalose import into peripheral tissues. Insect Biochemistry and Molecular Biology, 40(1), 30–37. https://doi.org/10.1016/j.ibmb.2009.12.006

51. Karouzou, M. V., Spyropoulos, Y., Iconomidou, V. A., Cornman, R. S., Hamodrakas, S. J., & Willis, J. H. (2007). Drosophila cuticular proteins with the R&R Consensus: Annotation and classification with a new tool for discriminating RR-1 and RR-2 sequences. Insect Biochemistry and Molecular Biology, 37(8), 754–760. https://doi.org/10.1016/j.ibmb.2007.03.007

52. Katoh, K. (2002). MAFFT: A novel method for rapid multiple sequence alignment based on fast Fourier transform. Nucleic Acids Research, 30(14), 3059–3066. https://doi.org/10.1093/nar/gkf436

53. Kemp, C., Mueller, S., Goto, A., Barbier, V., Paro, S., Bonnay, F., Dostert, C., Troxler, L., Hetru, C., Meignin, C., Pfeffer, S., Hoffmann, J. A., & Imler, J.-L. (2013). Broad RNA Interference–Mediated Antiviral Immunity and Virus-Specific Inducible Responses in *Drosophila*. The Journal of Immunology, 190(2), 650–658. https://doi.org/10.4049/jimmunol.1102486

54. Kongton, K., McCall, K., & Phongdara, A. (2014). Identification of gamma-interferon-inducible lysosomal thiol reductase (GILT) homologues in the fruit fly Drosophila melanogaster. Developmental & Comparative Immunology, 44(2), 389–396. https://doi.org/10.1016/j.dci.2014.01.007

55. Lake, C. M., Nielsen, R. J., Bonner, A. M., Eche, S., White-Brown, S., McKim, K. S., & Hawley, R. S. (2019). Narya, a RING finger domain-containing protein, is required for meiotic DNA double-strand break formation and crossover maturation in Drosophila melanogaster. PLOS Genetics, 15(1), e1007886. https://doi.org/10.1371/journal.pgen.1007886

56. Langmead, B., & Salzberg, S. L. (2012). Fast gapped-read alignment with Bowtie 2. Nature Methods, 9(4), 357–359. https://doi.org/10.1038/nmeth.1923

57. Lee, J. H., Cho, K. S., Lee, J., Yoo, J., Lee, J., & Chung, J. (2001). Diptericin-like protein: An immune response gene regulated by the anti-bacterial gene induction pathway in Drosophila. Gene, 271(2), 233–238. https://doi.org/10.1016/S0378-1119(01)00515-7

58. Lee, K.-A., Cho, K.-C., Kim, B., Jang, I.-H., Nam, K., Kwon, Y. E., Kim, M., Hyeon, D. Y., Hwang, D., Seol, J.-H., & Lee, W.-J. (2018). Inflammation-Modulated Metabolic Reprogramming Is Required for DUOX-Dependent Gut Immunity in Drosophila. Cell Host & Microbe, 23(3), 338–352.e5. https://doi.org/10.1016/j.chom.2018.01.011

59. Lemaitre, B., Reichhart, J.-M., & Hoffmann, J. A. (1997). *Drosophila* host defense: Differential induction of antimicrobial peptide genes after infection by various classes of microorganisms. Proceedings of the National Academy of Sciences, 94(26), 14614–14619. https://doi.org/10.1073/pnas.94.26.14614

60. Lerch, S., Zuber, R., Gehring, N., Wang, Y., Eckel, B., Klass, K.-D., Lehmann, F.-O., & Moussian, B. (2020). Resilin matrix distribution, variability and function in Drosophila. BMC Biology, 18(1), 195. https://doi.org/10.1186/s12915-020-00902-4

61. Levashina, E. A., Ohresser, S., Bulet, P., Reichhart, J.-M., Hetru, C., & Hoffmann, J. A. (1995). Metchnikowin, a Novel Immune-Inducible Proline-Rich Peptide from Drosophila with Antibacterial and Antifungal Properties. European Journal of Biochemistry, 233(2), 694–700. https://doi.org/10.1111/j.1432-1033.1995.694_2.x

62. Liao, Y., Smyth, G. K., & Shi, W. (2014). FeatureCounts: An efficient general purpose program for assigning sequence reads to genomic features. Bioinformatics. https://doi.org/10.1093/bioinformatics/btt656

63. Lin, Y., Golovnina, K., Chen, Z.-X., Lee, H. N., Negron, Y. L. S., Sultana, H., Oliver, B., & Harbison, S. T. (2016). Comparison of normalization and differential expression analyses using RNA-Seq data from 726 individual Drosophila melanogaster. BMC Genomics, 17(1), 28. https://doi.org/10.1186/s12864-015-2353-z

64. Linnemannstöns, K., Ripp, C., Honemann-Capito, M., Brechtel-Curth, K., Hedderich, M., & Wodarz, A. (2014). The PTK7-Related Transmembrane Proteins Off-track and Off-track 2 Are Co-receptors for Drosophila Wnt2 Required for Male Fertility. PLoS Genetics, 10(7), e1004443. https://doi.org/10.1371/journal.pgen.1004443

65. Litovchenko, M., Meireles-Filho, A. C. A., Frochaux, M. V., Bevers, R. P. J., Prunotto, A., Anduaga, A. M., Hollis, B., Gardeux, V., Braman, V. S., Russeil, J. M. C., Kadener, S., dal Peraro, M., & Deplancke, B. (2021). Extensive tissue-specific expression variation and novel regulators underlying circadian behavior. Science Advances, 7(5), eabc3781. https://doi.org/10.1126/sciadv.abc3781

66. Liu, Q., Kausar, S., Tang, Y., Huang, W., Tang, B., Abbas, M. N., & Dai, L. (2022). The Emerging Role of STING in Insect Innate Immune Responses and Pathogen Evasion Strategies. Frontiers in Immunology, 13, 874605. https://doi.org/10.3389/fimmu.2022.874605

67. Lollies, A. (2012). Wurst-Mediated Airway Clearance is Required for Postembryonic Development. Journal of Allergy & Therapy, 01(S7). https://doi.org/10.4172/2155-6121.S7-002

68. Longdon, B., Cao, C., Martinez, J., & Jiggins, F. M. (2013). Previous Exposure to an RNA Virus Does Not Protect against Subsequent Infection in Drosophila melanogaster. PLoS ONE, 8(9), e73833. https://doi.org/10.1371/journal.pone.0073833

69. Lu, Y., & Li, Z. (2015). Notch signaling downstream target E(spl)mbeta is dispensable for adult midgut homeostasis in Drosophila. Gene, 560(1), 89–95. https://doi.org/10.1016/j.gene.2015.01.053

70. Lung, O., & Wolfner, M. F. (2001). Identification and characterization of the major Drosophila melanogaster mating plug protein. Insect Biochemistry and Molecular Biology, 31(6–7), 543–551. https://doi.org/10.1016/S0965-1748(00)00154-5

71. Medd, N. C., Fellous, S., Waldron, F. M., Xuéreb, A., Nakai, M., Cross, J. V., & Obbard, D. J. (2018). The virome of Drosophila suzukii, an invasive pest of soft fruit. Virus Evolution, 4(1). https://doi.org/10.1093/ve/vey009

72. Meng, F. W., & Biteau, B. (2015). A Sox Transcription Factor Is a Critical Regulator of Adult Stem Cell Proliferation in the Drosophila Intestine. Cell Reports, 13(5), 906–914. https://doi.org/10.1016/j.celrep.2015.09.061

73. Meng, H., Yamashita, C., Shiba-Fukushima, K., Inoshita, T., Funayama, M., Sato, S., Hatta, T., Natsume, T., Umitsu, M., Takagi, J., Imai, Y., & Hattori, N. (2017). Loss of Parkinson’s disease-associated protein CHCHD2 affects mitochondrial crista structure and destabilizes cytochrome c. Nature Communications, 8(1), 15500. https://doi.org/10.1038/ncomms15500

74. Merkling, S. H., Overheul, G. J., van Mierlo, J. T., Arends, D., Gilissen, C., & van Rij, R. P. (2015). The heat shock response restricts virus infection in Drosophila. Scientific Reports, 5(1), 12758. https://doi.org/10.1038/srep12758

75. Minh, B. Q., Schmidt, H. A., Chernomor, O., Schrempf, D., Woodhams, M. D., von Haeseler, A., & Lanfear, R. (2020). IQ-TREE 2: New Models and Efficient Methods for Phylogenetic Inference in the Genomic Era. Molecular Biology and Evolution, 37(5), 1530–1534. https://doi.org/10.1093/molbev/msaa015

76. Mondotte, J. A., Gausson, V., Frangeul, L., Blanc, H., Lambrechts, L., & Saleh, M.-C. (2018). Immune priming and clearance of orally acquired RNA viruses in Drosophila. Nature Microbiology, 3(12), 1394–1403. https://doi.org/10.1038/s41564-018-0265-9

77. Mondotte, J. A., & Saleh, M.-C. (2018). Antiviral Immune Response and the Route of Infection in Drosophila melanogaster. In Advances in Virus Research (Vol. 100, pp. 247–278). Elsevier. https://doi.org/10.1016/bs.aivir.2017.10.006

78. Mussabekova, A., Daeffler, L., & Imler, J.-L. (2017). Innate and intrinsic antiviral immunity in Drosophila. Cellular and Molecular Life Sciences, 74(11), 2039– 2054. https://doi.org/10.1007/s00018-017-2453-9

79. Nayak, A., Berry, B., Tassetto, M., Kunitomi, M., Acevedo, A., Deng, C., Krutchinsky, A., Gross, J., Antoniewski, C., & Andino, R. (2010). Cricket paralysis virus antagonizes Argonaute 2 to modulate antiviral defense in Drosophila. Nature Structural & Molecular Biology, 17(5), 547–554. https://doi.org/10.1038/nsmb.1810

80. Okumura, T., Takeda, K., Kuchiki, M., Akaishi, M., Taniguchi, K., & Adachi-Yamada, T. (2016). GATAe regulates intestinal stem cell maintenance and differentiation in Drosophila adult midgut. Developmental Biology, 410(1), 24–35. https://doi.org/10.1016/j.ydbio.2015.12.017

81. O’Leary, N. A., Wright, M. W., Brister, J. R., Ciufo, S., Haddad, D., McVeigh, R., Rajput, B., Robbertse, B., Smith-White, B., Ako-Adjei, D., Astashyn, A., Badretdin, A., Bao, Y., Blinkova, O., Brover, V., Chetvernin, V., Choi, J., Cox, E., Ermolaeva, O., … Pruitt, K. D. (2016). Reference sequence (RefSeq) database at NCBI: Current status, taxonomic expansion, and functional annotation. Nucleic Acids Research. https://doi.org/10.1093/nar/gkv1189

82. Paine-Saunders, S., Fristrom, D., & Fristrom, J. W. (1990). The Drosophila IMP-E2 gene encodes an apically secreted protein expressed during imaginal disc morphogenesis. Developmental Biology, 140(2), 337–351. https://doi.org/10.1016/0012-1606(90)90084-V

83. Palmer, W. H., Joosten, J., Overheul, G. J., Jansen, P. W., Vermeulen, M., Obbard, D. J., & Van Rij, R. P. (2019). Induction and Suppression of NF-κB Signalling by a DNA Virus of *Drosophila*. Journal of Virology, 93(3), e01443–18. https://doi.org/10.1128/JVI.01443-18

84. Palmer, W. H., Medd, N. C., Beard, P. M., & Obbard, D. J. (2018). Isolation of a natural DNA virus of Drosophila melanogaster, and characterisation of host resistance and immune responses. PLOS Pathogens, 14(6), e1007050. https://doi.org/10.1371/journal.ppat.1007050

85. Qu, W., Gurdziel, K., Pique-Regi, R., & Ruden, D. M. (2017). Identification of Splicing Quantitative Trait Loci (sQTL) in Drosophila melanogaster with Developmental Lead (Pb2+) Exposure. Frontiers in Genetics, 8, 145. https://doi.org/10.3389/fgene.2017.00145

86. R Core Team. (2021). R: A language and environment for statistical computing. R Foundation for Statistical Computing, Vienna, Austria. https://www.R-project.org/

87. Ross, J., Jiang, H., Kanost, M. R., & Wang, Y. (2003). Serine proteases and their homologs in the Drosophila melanogaster genome: An initial analysis of sequence conservation and phylogenetic relationships. Gene, 304, 117–131. https://doi.org/10.1016/S0378-1119(02)01187-3

88. Schnakenberg, S. L., Matias, W. R., & Siegal, M. L. (2011). Sperm-Storage Defects and Live Birth in Drosophila Females Lacking Spermathecal Secretory Cells. PLoS Biology, 9(11), e1001192. https://doi.org/10.1371/journal.pbio.1001192

89. Shelly, S., Lukinova, N., Bambina, S., Berman, A., & Cherry, S. (2009). Autophagy Is an Essential Component of Drosophila Immunity against Vesicular Stomatitis Virus. Immunity, 30(4), 588–598. https://doi.org/10.1016/j.immuni.2009.02.009

90. Shi, M., White, V. L., Schlub, T., Eden, J.-S., Hoffmann, A. A., & Holmes, E. C. (2018). No detectable effect of *Wolbachia w* Mel on the prevalence and abundance of the RNA virome of *Drosophila melanogaster*. Proceedings of the Royal Society B: Biological Sciences, 285(1883), 20181165. https://doi.org/10.1098/rspb.2018.1165

91. Shukla, A. K., Spurrier, J., Kuzina, I., & Giniger, E. (2019). Hyperactive Innate Immunity Causes Degeneration of Dopamine Neurons upon Altering Activity of Cdk5. Cell Reports, 26(1), 131–144.e4. https://doi.org/10.1016/j.celrep.2018.12.025

92. Sitnik, J. L., Gligorov, D., Maeda, R. K., Karch, F., & Wolfner, M. F. (2016). The Female Post-Mating Response Requires Genes Expressed in the Secondary Cells of the Male Accessory Gland in *Drosophila melanogaster*. Genetics, 202(3), 1029–1041. https://doi.org/10.1534/genetics.115.181644

93. Subramanian, M., Metya, S. K., Sadaf, S., Kumar, S., Schwudke, D., & Hasan, G. (2013). Altered lipid homeostasis in *Drosophila* InsP3 receptor mutants leads to obesity and hyperphagia. Disease Models & Mechanisms, dmm.010017. https://doi.org/10.1242/dmm.010017

94. Syed, Z. A., Härd, T., Uv, A., & Van Dijk-Härd, I. F. (2008). A Potential Role for Drosophila Mucins in Development and Physiology. PLoS ONE, 3(8), e3041. https://doi.org/10.1371/journal.pone.0003041

95. Tafesh-Edwards, G., & Eleftherianos, I. (2020). Drosophila immunity against natural and nonnatural viral pathogens. Virology, 540, 165–171. https://doi.org/10.1016/j.virol.2019.12.001

96. Teninges, D., Ohanessian, A., Richard-Molard, C., & Contamine, D. (1979). Isolation and Biological Properties of Drosophila X Virus. Journal of General Virology, 42(2), 241–254. https://doi.org/10.1099/0022-1317-42-2-241

97. Tian, C., Gao, B., Rodriguez, M. D. C., Lanz-Mendoza, H., Ma, B., & Zhu, S. (2008). Gene expression, antiparasitic activity, and functional evolution of the drosomycin family. Molecular Immunology, 45(15), 3909–3916. https://doi.org/10.1016/j.molimm.2008.06.025

98. Tootle, T. L., Williams, D., Hubb, A., Frederick, R., & Spradling, A. (2011). Drosophila Eggshell Production: Identification of New Genes and Coordination by Pxt. PLoS ONE, 6(5), e19943. https://doi.org/10.1371/journal.pone.0019943

99. van Mierlo, J. T., Overheul, G. J., Obadia, B., van Cleef, K. W. R., Webster, C. L., Saleh, M.-C., Obbard, D. J., & van Rij, R. P. (2014). Novel Drosophila Viruses Encode Host-Specific Suppressors of RNAi. PLoS Pathogens, 10(7), e1004256. https://doi.org/10.1371/journal.ppat.1004256

100. Verleyen, P., Baggerman, G., D’Hertog, W., Vierstraete, E., Husson, S. J., & Schoofs, L. (2006). Identification of new immune induced molecules in the haemolymph of Drosophila melanogaster by 2D-nanoLC MS/MS. Journal of Insect Physiology, 52(4), 379–388. https://doi.org/10.1016/j.jinsphys.2005.12.007

101. Vig, M., Peinelt, C., Beck, A., Koomoa, D. L., Rabah, D., Koblan-Huberson, M., Kraft, S., Turner, H., Fleig, A., Penner, R., & Kinet, J.-P. (2006). CRACM1 Is a Plasma Membrane Protein Essential for Store-Operated Ca ^2+^ Entry. Science, 312(5777), 1220–1223. https://doi.org/10.1126/science.1127883

102. Vorbrüggen, G., Constien, R., Zilian, O., Wimmer, E. A., Dowe, G., Taubert, H., Noll, M., & Jäckle, H. (1997). Embryonic expression and characterization of a Ptx1 homolog in Drosophila. Mechanisms of Development, 68(1–2), 139–147. https://doi.org/10.1016/S0925-4773(97)00139-1

103. Voßfeldt, H., Butzlaff, M., Prüßing, K., Ní Chárthaigh, R.-A., Karsten, P., Lankes, A., Hamm, S., Simons, M., Adryan, B., Schulz, J. B., & Voigt, A. (2012). Large-Scale Screen for Modifiers of Ataxin-3-Derived Polyglutamine-Induced Toxicity in Drosophila. PLoS ONE, 7(11), e47452. https://doi.org/10.1371/journal.pone.0047452

104. Wallace, M. A., Coffman, K. A., Gilbert, C., Ravindran, S., Albery, G. F., Abbott, J., Argyridou, E., Bellosta, P., Betancourt, A. J., Colinet, H., Eric, K., Glaser-Schmitt, A., Grath, S., Jelic, M., Kankare, M., Kozeretska, I., Loeschcke, V., Montchamp-Moreau, C., Ometto, L., … Obbard, D. J. (2021). The discovery, distribution, and diversity of DNA viruses associated with *Drosophila melanogaster* in Europe. Virus Evolution, 7(1), veab031. https://doi.org/10.1093/ve/veab031

105. Wang, L., Jahren, N., Vargas, M. L., Andersen, E. F., Benes, J., Zhang, J., Miller, E. L., Jones, R. S., & Simon, J. A. (2006). Alternative ESC and ESC-Like Subunits of a Polycomb Group Histone Methyltransferase Complex Are Differentially Deployed during *Drosophila* Development. Molecular and Cellular Biology, 26(7), 2637–2647. https://doi.org/10.1128/MCB.26.7.2637-2647.2006

106. Wang, X.-H., Aliyari, R., Li, W.-X., Li, H.-W., Kim, K., Carthew, R., Atkinson, P., & Ding, S.-W. (2006). RNA Interference Directs Innate Immunity Against Viruses in Adult *Drosophila*. Science, 312(5772), 452–454. https://doi.org/10.1126/science.1125694

107. Watkins, P. A., Maiguel, D., Jia, Z., & Pevsner, J. (2007). Evidence for 26 distinct acyl-coenzyme A synthetase genes in the human genome. Journal of Lipid Research, 48(12), 2736–2750. https://doi.org/10.1194/jlr.M700378-JLR200

108. Webster, C. L., Longdon, B., Lewis, S. H., & Obbard, D. J. (2016). Twenty-Five New Viruses Associated with the Drosophilidae (Diptera). Evolutionary Bioinformatics, 12s2, EBO.S39454. https://doi.org/10.4137/EBO.S39454

109. Webster, C. L., Waldron, F. M., Robertson, S., Crowson, D., Ferrari, G., Quintana, J. F., Brouqui, J.-M., Bayne, E. H., Longdon, B., Buck, A. H., Lazzaro, B. P., Akorli, J., Haddrill, P. R., & Obbard, D. J. (2015). The Discovery, Distribution, and Evolution of Viruses Associated with Drosophila melanogaster. PLOS Biology, 13(7), e1002210. https://doi.org/10.1371/journal.pbio.1002210

110. Wijffels, G., Eisemann, C., Riding, G., Pearson, R., Jones, A., Willadsen, P., & Tellam, R. (2001). A Novel Family of Chitin-binding Proteins from Insect Type 2 Peritrophic Matrix. Journal of Biological Chemistry, 276(18), 15527–15536. https://doi.org/10.1074/jbc.M009393200

111. Wolfe, J., Akam, M. E., & Roberts, D. B. (1977). Biochemical and Immunological Studies on Larval Serum Protein 1, the Major Haemolymph Protein of Drosophila melanogaster Third-Instar Larvae. European Journal of Biochemistry, 79(1), 47–53. https://doi.org/10.1111/j.1432-1033.1977.tb11782.x

112. Wong, Z. S., Brownlie, J. C., & Johnson, K. N. (2016). Impact of ERK activation on fly survival and Wolbachia-mediated protection during virus infection. Journal of General Virology, 97(6), 1446–1452. https://doi.org/10.1099/jgv.0.000456

113. Zambon, R. A., Nandakumar, M., Vakharia, V. N., & Wu, L. P. (2005). The Toll pathway is important for an antiviral response in *Drosophila*. Proceedings of the National Academy of Sciences, 102(20), 7257–7262. https://doi.org/10.1073/pnas.0409181102

114. Zhang, L., Xu, W., Gao, X., Li, W., Qi, S., Guo, D., Ajayi, O. E., Ding, S.-W., & Wu, Q. (2020). LncRNA Sensing of a Viral Suppressor of RNAi Activates Non-canonical Innate Immune Signaling in Drosophila. Cell Host & Microbe, 27(1), 115–128.e8. https://doi.org/10.1016/j.chom.2019.12.006

115. Zhong, W., McClure, C. D., Evans, C. R., Mlynski, D. T., Immonen, E., Ritchie, M. G., & Priest, N. K. (2013). Immune anticipation of mating in *Drosophila*: *Turandot M* promotes immunity against sexually transmitted fungal infections. Proceedings of the Royal Society B: Biological Sciences, 280(1773), 20132018. https://doi.org/10.1098/rspb.2013.2018

116. Zhou, F., Qiang, K. M., & Beckingham, K. M. (2016). Failure to Burrow and Tunnel Reveals Roles for jim lovell in the Growth and Endoreplication of the Drosophila Larval Tracheae. PLOS ONE, 11(8), e0160233. https://doi.org/10.1371/journal.pone.0160233

117. Zhu, F., Ding, H., & Zhu, B. (2013). Transcriptional profiling of Drosophila S2 cells in early response to Drosophila C virus. Virology Journal, 10(1), 210. https://doi.org/10.1186/1743-422X-10-210

118. Zuber, R., Wang, Y., Gehring, N., Bartoszewski, S., & Moussian, B. (2020). Tweedle proteins form extracellular two-dimensional structures defining body and cell shape in *Drosophila melanogaster*. Open Biology, 10(12), 200214. https://doi.org/10.1098/rsob.200214

