## Supplementary material for "Viruses in laboratory *Drosophila* and their impact on host gene expression": Document S1

**Estimating Barcode Switching Rates**

Barcode switching or ‘hopping’ can result in the misassignment of reads among libraries (Griffiths et al., 2018). Although the impact is likely to be extremely small for analysis of host gene expression, where no gene of interest constitutes the bulk of the library, it could easily cause a sample to be misclassified as ‘infected’ with a virus that is in reality absent but present at very high copy-number in another sample. Note that such barcode switching would not induce false positives in our analysis, but by blurring the lines between ‘infected’ and ‘uninfected’, it may reduce power. To mitigate this, we attempted to estimate the rate of barcode switching in these public datasets and use this to inform a threshold number of virus reads for classification as ‘infected’.

First, the inter-sex barcode switching per flow cell lane was estimated as the total proportion of sex-specific reads that were wrongly assigned to flies of the opposites (**Equation 1**). Second, the intra-sex barcode switching was inferred as the proportion of sex-specific reads that were likely wrongly assigned to flies of the same sex, based on the product of intersex barcode switching and the proportion of males and female flies in the flow cell lane (**Equation 2**). Finally, the total proportion of barcode switching per dataset was estimated as the sum of intersex and intersex barcode switching (**Equation 3**). For flow cell lanes that had only one sex, I used the average proportion of barcode switching from other lanes in the same dataset in which both sexes were represented. If the entire dataset had only one sex, I used the estimated barcode switching from datasets that were sequenced using similar platforms. The number of virus reads that were affected by barcode switching in each library was then estimated from the average proportion of barcode switching. The presence of a virus in a library was confirmed if the number of reads that mapped to the virus was greater than 150 and was over twice the error estimated for barcode switching. A threshold of 150 reads was selected since this threshold reduces the inconsistency of duplicate samples (**Figure S1**).

***Equation 1*** $Intersex_{BS} = \frac{M}{TF} + \frac{F}{TM}$

***Equation 2*** $Intrasex_{BS} = \frac{M}{TF} \times \frac{NM}{NFly} + \frac{F}{TM} \times \frac{NF}{NFly}$

***Equation 3*** $Proportion_{BS} = Intersex_{BS} + Intrasex_{BS}$

**Equations 1 to 3 key**

**M** = Sum of male-specific reads that were wrongly assigned to females

**F** = Sum of female-specific reads that were wrongly assigned to males

**TM** = Sum of reads that were assigned to male-specific genes

**TF** = Sum of reads that were assigned to female-specific genes

**NM**= Number of males

**NM**= Number of females

**NFly**= Number of libraries
