## Supplementary material for "Viruses in laboratory *Drosophila* and their impact on host gene expression": Document S2

#### Nora virus

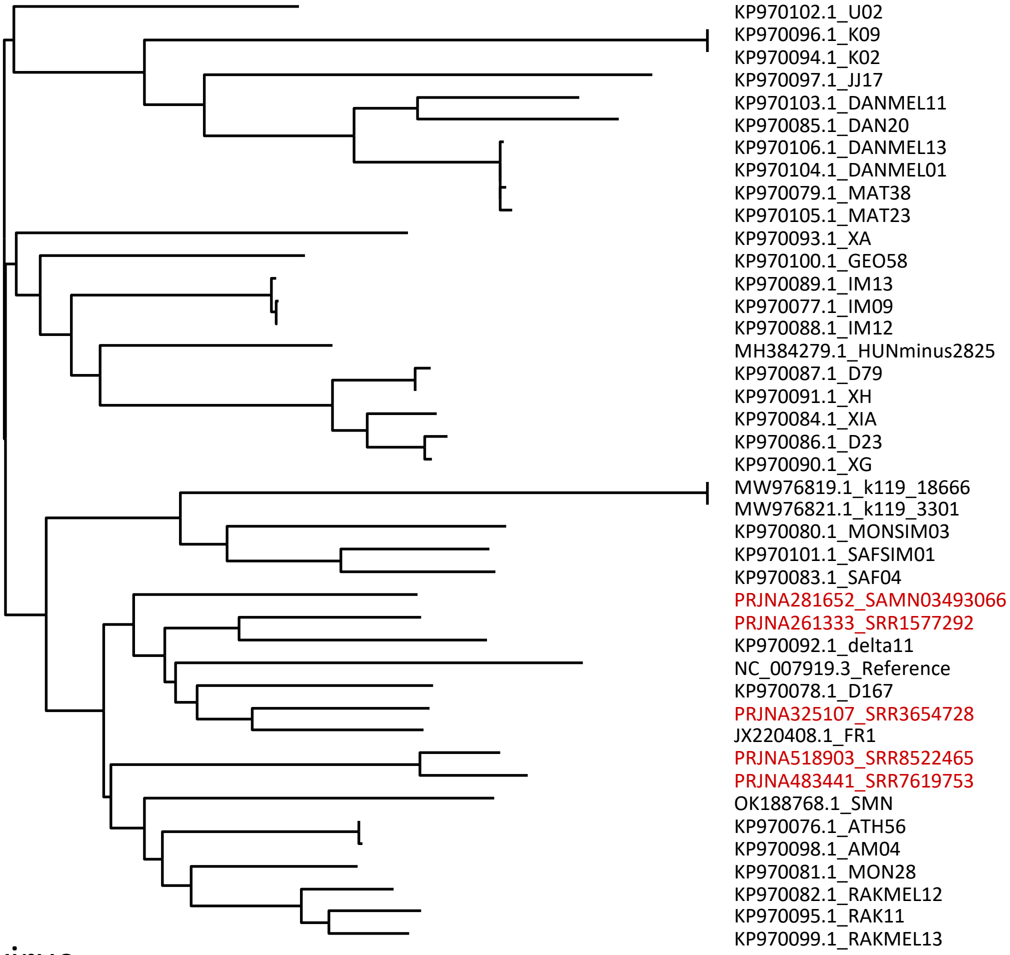


#### Thika virus

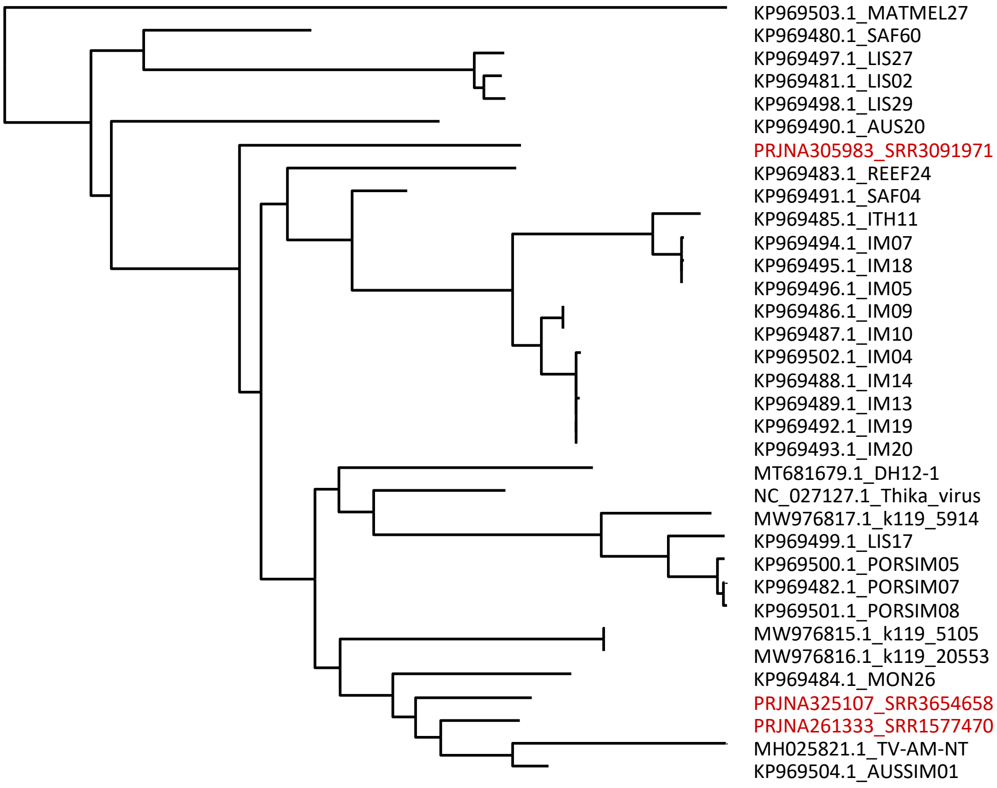

#### Drosophila A virus

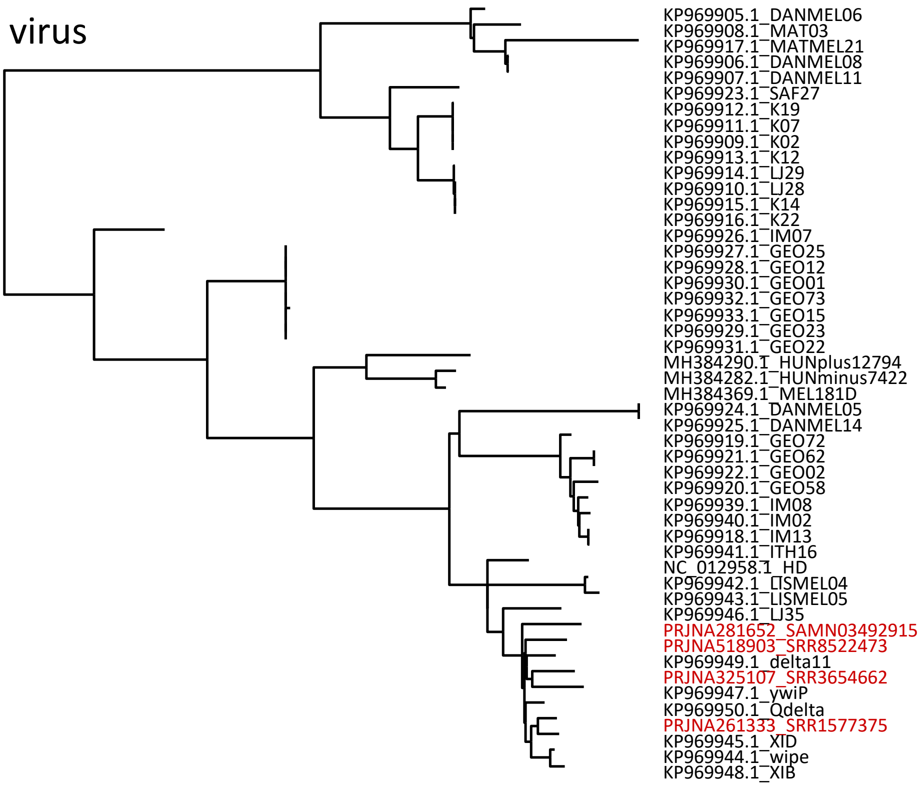


#### Drosophila C virus

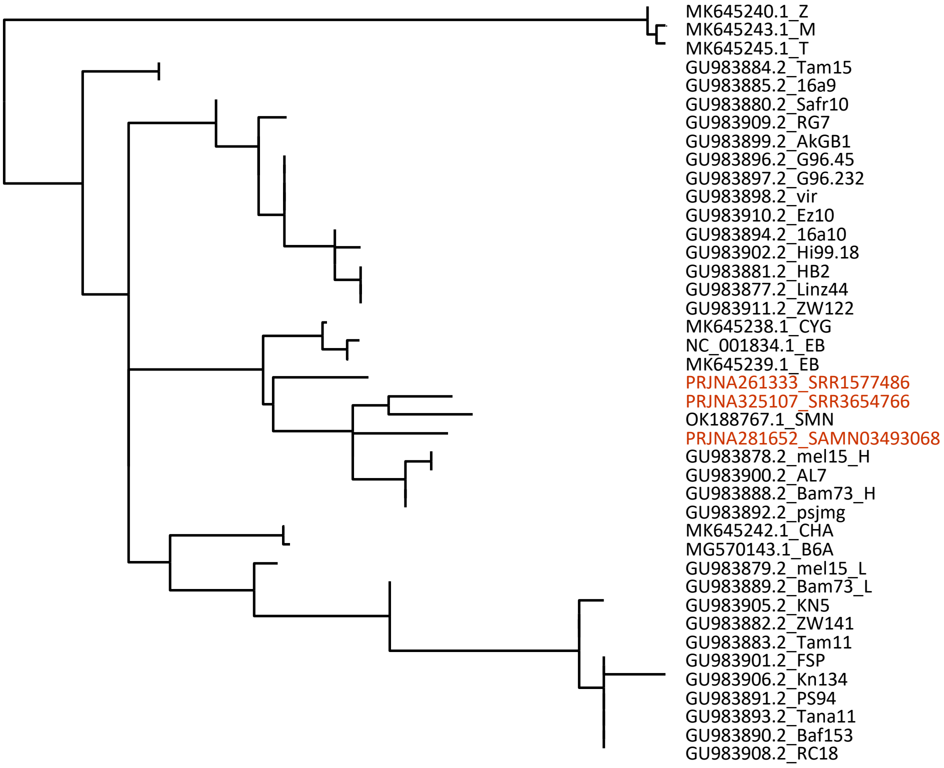

### Craigies Hill virus (Segment 1)

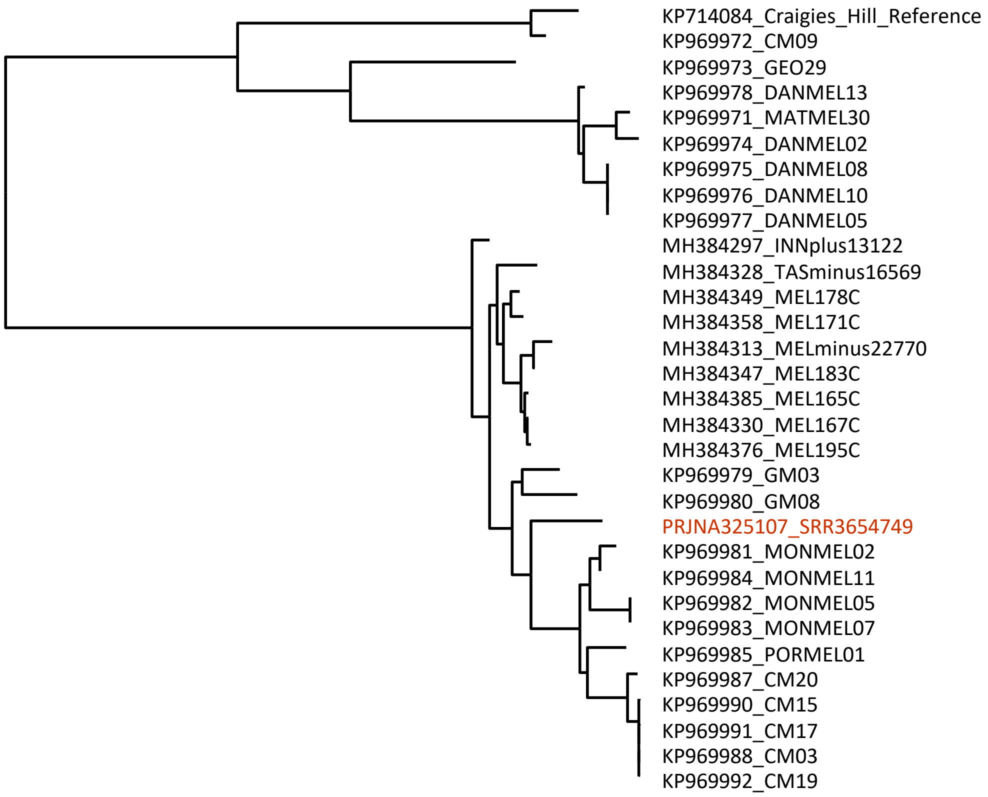


### Bloomfield virus (Segment 1)

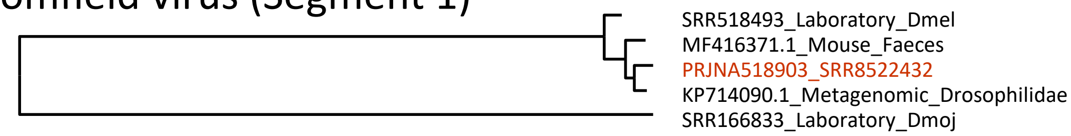


### Brandeis virus

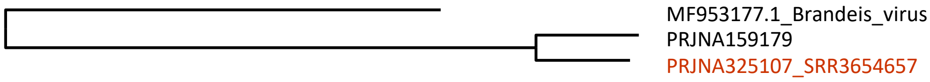

#### Galbut virus (Segment 2)

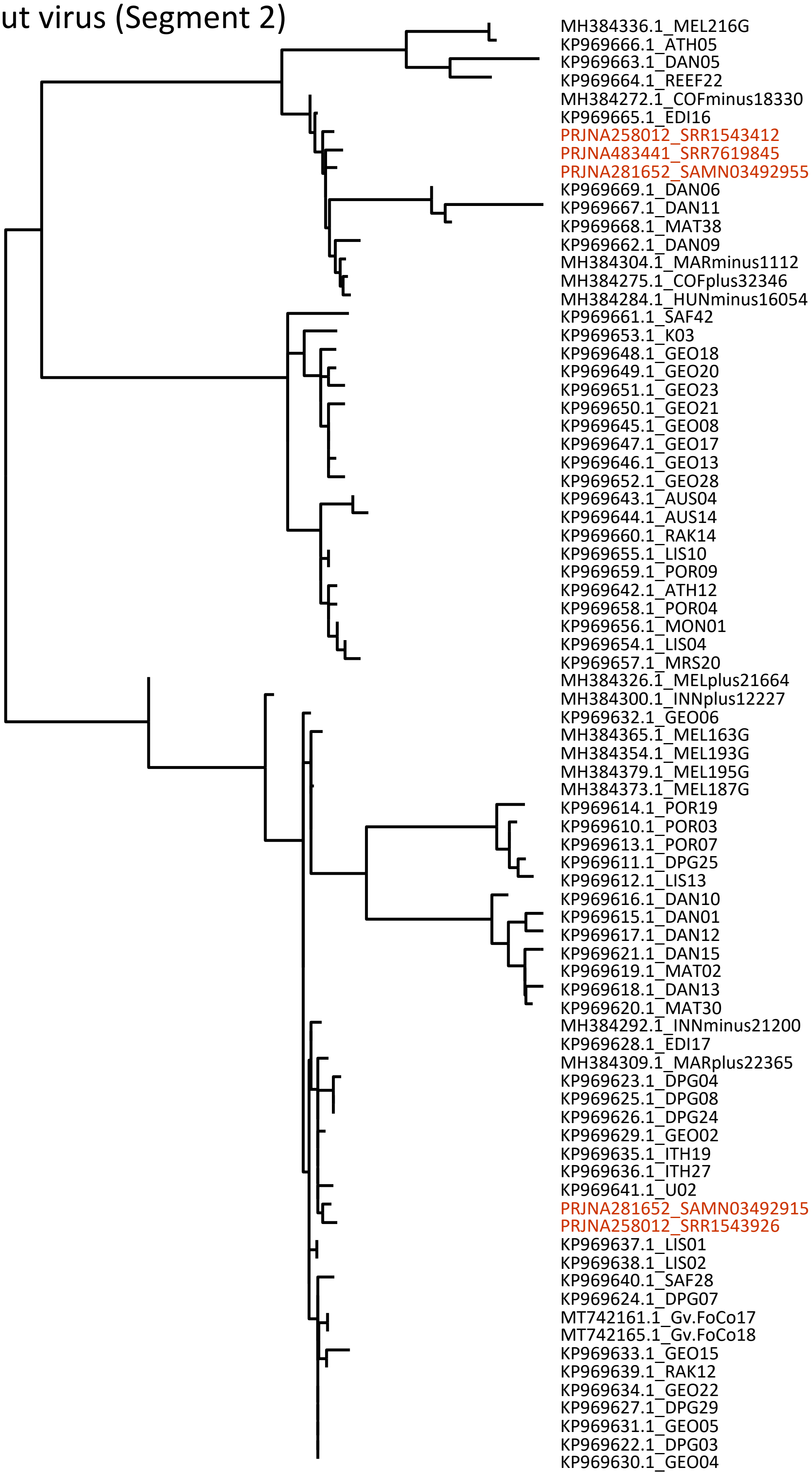

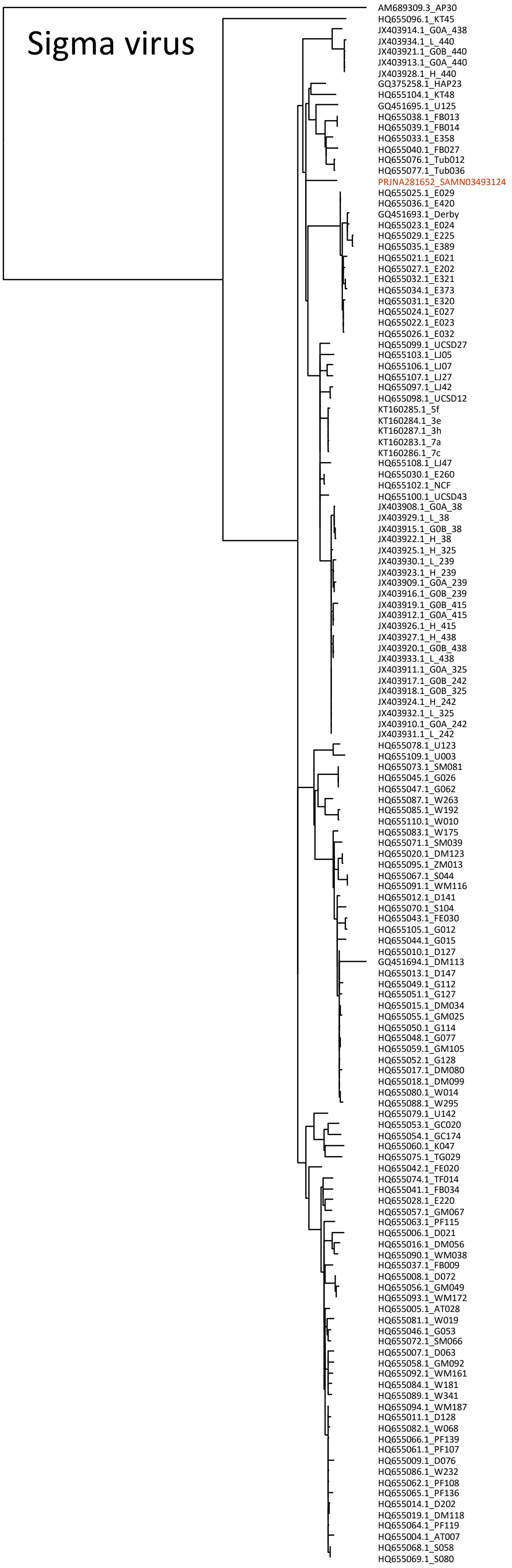
Sigma virus
AM689309.3_AP30
HQ655096.1_KT45
JX403914.1_G0A_438
JX403934.1_L_440
JX403921.1_G0B_440
JX403913.1_G0A_440
JX403928.1_H_440
GQ375258.1_HAP23
HQ655104.1_KT48
GQ451695.1_U125
HQ655038.1_FB013
HQ655039.1_FB014
HQ655033.1_E358
HQ655040.1_FB027
HQ655076.1_Tub012
HQ655077.1_Tub036
PRJNA281652_SAMN03493124
HQ655025.1_E029
HQ655036.1_E420
GQ451693.1_Derby
HQ655023.1_E024
HQ655029.1_E225
HQ655035.1_E389
HQ655021.1_E021
HQ655027.1_E202
HQ655032.1_E321
HQ655034.1_E373
HQ655031.1_E320
HQ655024.1_E027
HQ655022.1_E023
HQ655026.1_E032
HQ655099.1_UCSD27
HQ655103.1_LJ05
HQ655106.1_LJ07
HQ655107.1_LJ27
HQ655097.1_LJ42
HQ655098.1_UCSD12
KT160285.1_5f
KT160284.1_3e
KT160287.1_3h
KT160283.1_7a
KT160286.1_7c
HQ655108.1_LJ47
HQ655030.1_E260
HQ655102.1_NCF
HQ655100.1_UCSD43
JX403908.1_G0A_38
JX403929.1_L_38
JX403915.1_G0B_38
JX403922.1_H_38
JX403925.1_H_325
JX403930.1_L_239
JX403923.1_H_239
JX403909.1_G0A_239
JX403916.1_G0B_239
JX403919.1_G0B_415
JX403912.1_G0A_415
JX403926.1_H_415
JX403927.1_H_438
JX403920.1_G0B_438
JX403933.1_L_438
JX403911.1_G0A_325
JX403917.1_G0B_242
JX403918.1_G0B_325
JX403924.1_H_242
JX403932.1_L_325
JX403910.1_G0A_242
JX403931.1_L_242
HQ655078.1_U123
HQ655109.1_U003
HQ655073.1_SM081
HQ655045.1_G026
HQ655047.1_G062
HQ655087.1_W263
HQ655085.1_W192
HQ655110.1_W010
HQ655083.1_W175
HQ655071.1_SM039
HQ655020.1_DM123
HQ655095.1_ZM013
HQ655067.1_S044
HQ655091.1_WM116
HQ655012.1_D141
HQ655070.1_S104
HQ655043.1_FE030
HQ655105.1_G012
HQ655044.1_G015
HQ655010.1_D127
GQ451694.1_DM113
HQ655013.1_D147
HQ655049.1_G112
HQ655051.1_G127
HQ655015.1_DM034
HQ655055.1_GM025
HQ655050.1_G114
HQ655048.1_G077
HQ655059.1_GM105
HQ655052.1_G128
HQ655017.1_DM080
HQ655018.1_DM099
HQ655080.1_W014
HQ655088.1_W295
HQ655079.1_U142
HQ655053.1_GC020
HQ655054.1_GC174
HQ655060.1_K047
HQ655075.1_TG029
HQ655042.1_FE020
HQ655074.1_TF014
HQ655041.1_FB034
HQ655028.1_E220
HQ655057.1_GM067
HQ655063.1_PF115
HQ655006.1_D021
HQ655016.1_DM056
HQ655090.1_WM038
HQ655037.1_FB009
HQ655008.1_D072
HQ655056.1_GM049
HQ655093.1_WM172
HQ655005.1_AT028
HQ655081.1_W019
HQ655046.1_G053
HQ655072.1_SM066
HQ655007.1_D063
HQ655058.1_GM092
HQ655092.1_WM161
HQ655084.1_W181
HQ655089.1_W341
HQ655094.1_WM187
HQ655011.1_D128
HQ655082.1_W068
HQ655066.1_PF139
HQ655061.1_PF107
HQ655009.1_D076
HQ655086.1_W232
HQ655062.1_PF108
HQ655065.1_PF136
HQ655014.1_D202
HQ655019.1_DM118
HQ655064.1_PF119
HQ655004.1_AT007
HQ655068.1_S058
HQ655069.1_S080

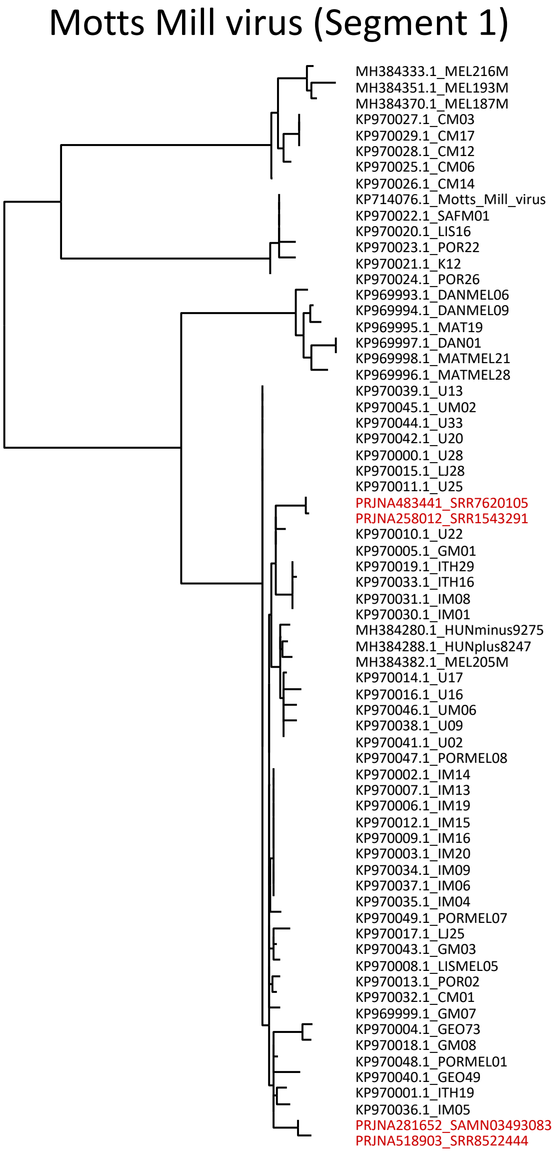
Motts Mill virus (Segment 1)
MH384333.1_MEL216M
MH384351.1_MEL193M
MH384370.1_MEL187M
KP970027.1_CM03
KP970029.1_CM17
KP970028.1_CM12
KP970025.1_CM06
KP970026.1_CM14
KP714076.1_Motts_Mill_virus
KP970022.1_SAFM01
KP970020.1_LIS16
KP970023.1_POR22
KP970021.1_K12
KP970024.1_POR26
KP969993.1_DANMEL06
KP969994.1_DANMEL09
KP969995.1_MAT19
KP969997.1_DAN01
KP969998.1_MATMEL21
KP969996.1_MATMEL28
KP970039.1_U13
KP970045.1_UM02
KP970044.1_U33
KP970042.1_U20
KP970000.1_U28
KP970015.1_LJ28
KP970011.1_U25
PRJNA483441_SRR7620105
PRJNA258012_SRR1543291
KP970010.1_U22
KP970005.1_GM01
KP970019.1_ITH29
KP970033.1_ITH16
KP970031.1_IM08
KP970030.1_IM01
MH384280.1_HUNminus9275
MH384288.1_HUNplus8247
MH384382.1_MEL205M
KP970014.1_U17
KP970016.1_U16
KP970046.1_UM06
KP970038.1_U09
KP970041.1_U02
KP970047.1_PORMEL08
KP970002.1_IM14
KP970007.1_IM13
KP970006.1_IM19
KP970012.1_IM15
KP970009.1_IM16
KP970003.1_IM20
KP970034.1_IM09
KP970037.1_IM06
KP970035.1_IM04
KP970049.1_PORMEL07
KP970017.1_LJ25
KP970043.1_GM03
KP970008.1_LISMEL05
KP970013.1_POR02
KP970032.1_CM01
KP969999.1_GM07
KP970004.1_GEO73
KP970018.1_GM08
KP970048.1_PORMEL01
KP970040.1_GEO49
KP970001.1_ITH19
KP970036.1_IM05
PRJNA281652_SAMN03493083
PRJNA518903_SRR8522444
