## Supplementary material for "Viruses in laboratory *Drosophila* and their impact on host gene expression": Figure S1

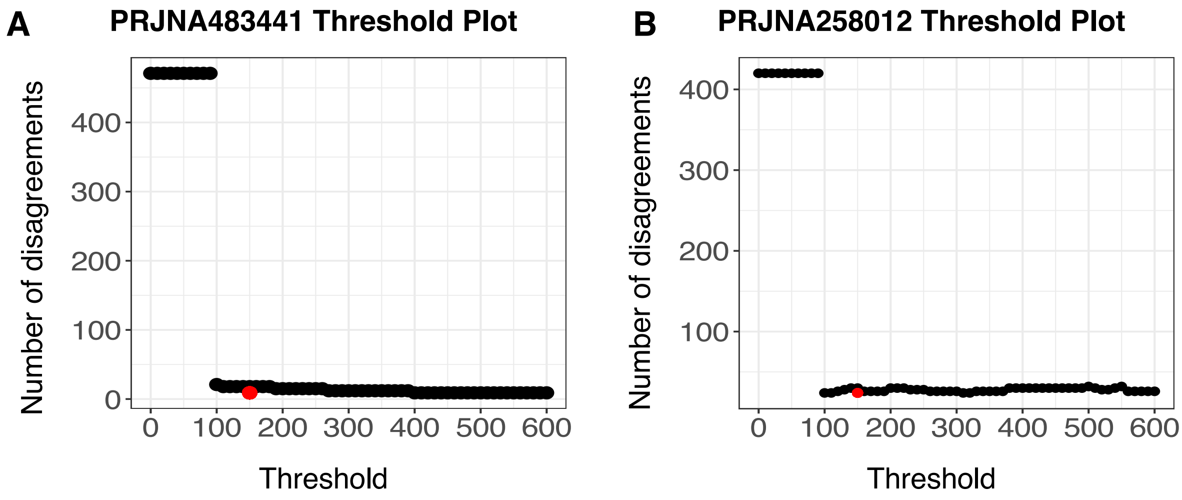


**Figure S1 Best threshold to estimate viral presence**

The datasets PRJNA483441 (**A**) and PRJNA258012 (**B**) have duplicate libraries. We compared the number of duplicates where there were disagreements in virus infection status when different threshold values from 0 to 600 were used i.e., the number of duplicates where one has present and the other has absent for a particular virus. The threshold at 150 is coloured red.
