## Supplementary material for "Viruses in laboratory *Drosophila* and their impact on host gene expression": Figure S2

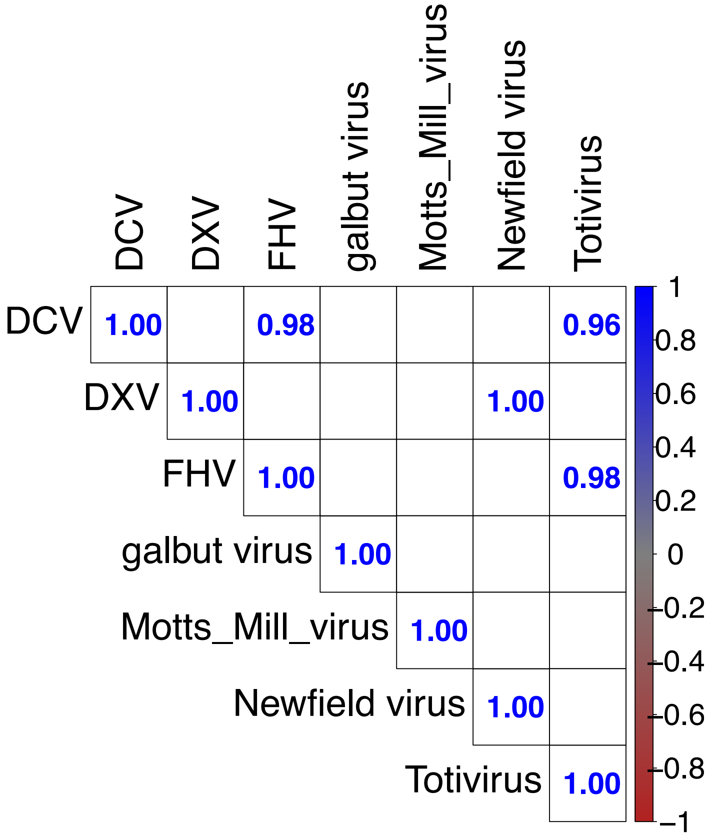


**Figure S2 Correlations between normalised reads in PRJNA258012**

The figure shows significant ($p < 0.001$) r values obtained via Spearman correlation between the normalised virus reads (RPKM) in samples in the dataset PRJNA258012. Blank squares have non-significant correlations.
