## Supplementary figures and images for "Viruses in laboratory *Drosophila* and their impact on host gene expression"

### Figure S3

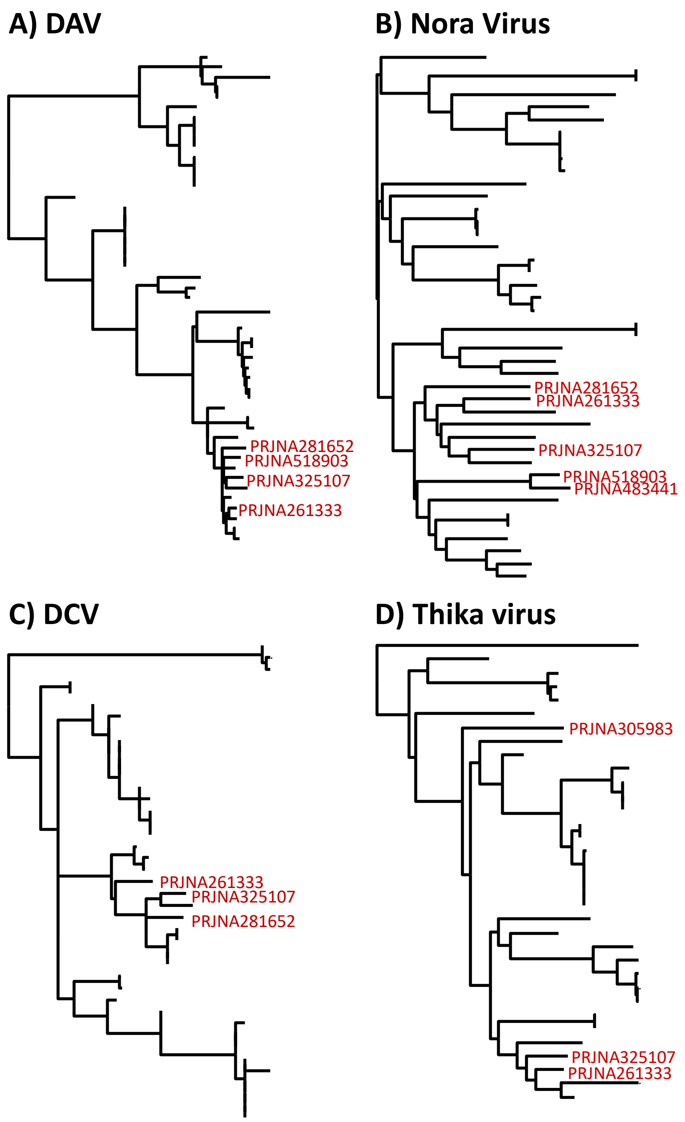
