## Supplementary material for "Viruses in laboratory *Drosophila* and their impact on host gene expression": Figure S4

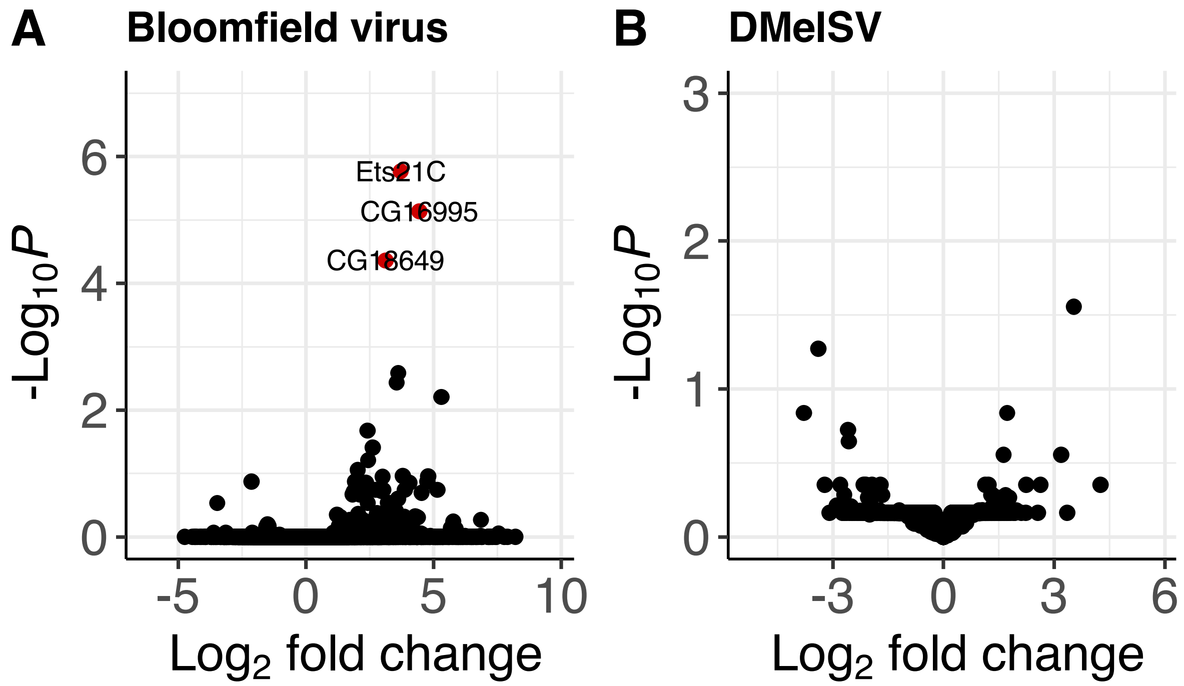


**Figure S3 Effects of viral infection on host gene expression**

Using uninfected females as baseline, the volcano plots show the effects of viral infection. The virus names and total number of variables (genes) used in each expression analysis is shown. None of the genes had significant changes in expression.
